# An engineered chromatin protein with enhanced preferential binding of H3K27me3 over H3K9me3

**DOI:** 10.1101/2024.10.02.616304

**Authors:** Kierra A. Franklin, J. Harrison Priode, Arefeh Abbasi, Ivan Kordic, Joongho Sohn, Paige Steppe, Ahmet Coskun, Esther W. Gomez, Karmella A. Haynes

## Abstract

Histone post-translational modifications (PTMs) in chromatin modulate enhancer function, gene expression, and cellular differentiation. Pockets within chromatin reader-effector proteins bind specific histone PTMs, and have informed the design of artificial probes. Histone binding domains (HBDs) interact via multiple contacts that may support greater specificity than antibodies, but are difficult to study because of their weak affinities *in vitro*. We used a “Cell-Free Histone-Binding Immunoassay” (CHIA) where interactions between cell-free-expressed engineered HBD proteins and immobilized histone peptides are measured in an ELISA system. Previously reported K33E and Q9D variants of the chromobox 7 (CBX7) HBD bound with high affinity to both H3K27me3 and H3K9me3. However in CBX8, the K33E substitution enhanced binding to K27me3 with minimal K9me3 binding. Furthermore, hydrophobic substitutions in the CBX8 hydrophobic clasp (V10, L49) supported affinity and specificity for K27me3. We demonstrate the utility of a new CBX8 variant as a H3K27me3-specific live cell chromatin probe.

## Introduction

Chromatin is a complex of DNA and proteins that enables a wide range of essential processes such as genome organization and remodeling, transcriptional regulation, and DNA repair in eukaryotes. Histone post-translational modifications (PTMs), such as acetylation, methylation, and phosphorylation, are covalent modifications concentrated on the unstructured N-terminal regions of histones that modulate these essential processes by acting as ligands for chromatin proteins. Lysine methylation is a critical determinant of cell state and function and its aberrant expression has been indicated as a key contributor to disease (reviewed in Franklin et al^1^). Thus, the ability to reliably detect histone PTM and their interactions with other proteins is necessary to better understand disease pathology as well as chromatin processes that underlie gene regulation.

Currently, mapping of histone PTMs is still largely achieved through antibody detection. However, anti-histone PTM antibodies are notoriously unreliable due to frequent cross-reactivity with off-target histone modifications and substantial lot-to-lot variability^2,3^. These limitations make it difficult to tease apart the roles of distinct histone PTMs in chromatin regulation. Therefore, scientists have developed histone PTM-binding domains (HBDs) to expand the arsenal of detection probes beyond antibodies. A family of naturally occurring peptide folds known as epigenetic “readers” are gaining popularity as research tools for their ability to recognize histone acetylation, methylation, phosphorylation, and DNA methylation^1,4^. The use of HBDs as antibody alternatives holds great promise considering their small size and ability to discriminate various histone PTMs. Wild-type and engineered HBDs have been used for biochemical assays, live-cell histone PTM tracking, and chromatin mapping^5,6^. Improving HBD binding affinity is an important step to achieve robust PTM recognition at low protein concentrations.

To generate high-affinity readers of lysine methylation, scientists have used the chromobox (CBX) family of proteins from multicellular organisms as the basis of rational design and protein variant library screens^7,8^. Chromobox proteins are characterized by a conserved N-terminal domain that contains a pocket of aromatic residues, typically phenylalanine (F), tryptophan (W), and/or tyrosine (Y), that enclose the target histone methyl-lysine. A chromobox subgroup that includes CBX1, 3, and 5 contain an N-terminal chromodomain (CD) that preferentially binds histone PTM H3K9me3. Recent efforts have used structure-based computational modeling to predict mutations that improve CBX1 affinity to H3K9me3, but less attention has been paid to engineering binders for H3K27me3^9,10^.

The CBX2, 4, 6, 7 and 8 chromobox proteins contain an N-terminal polycomb chromodomain (PCD) that interacts with histone PTM H3K27me3, followed by a disordered positively-charged domain of variable length that supports chromatin compaction, and a C box domain that supports the incorporation of the CBX protein into canonical polycomb repressive complex 1 (cPRC1)^11^. Isolated PCDs from CBX8 show weak binding affinities for H3K27me3 (Kd 5 to >500 μM), but are highly specific^12,13,14^. We previously showed that the affinity can be enhanced by adding a second N-terminal PCD. Recently Veggiani et al. identified a high-affinity CBX7 variant, CBX7.VD, in a phage display screen for strong monovalent binders of H3K27me3^15^. However, this variant showed strong affinity for both H3K27me3 and H3K9me3. Discriminating these two PTMs may be difficult because both methyl-lysines appear in a ‘ARKS’ motif on the histone 3 tail. Specific structural features that enable the discrimination of the two marks have been the subject of intense study^12,13,16,17^. Therefore we set out to gain deeper insights into PCD structure and function, as well as develop an improved probe for H3K27me3, by enhancing the affinity of a single PCD.

Here, we report substitutions of amino acids upstream of the aromatic H3K27me3-binding pocket of the CBX8 polycomb chromodomain that enhance its affinity for H3K27me3. We used a library of 3072 CBX8 PCD variants to build new bivalent proteins with a customized Golden Gate assembly method^18^. We show that our Cell-Free Histone-Binding Immunoassay (CHIA) is sensitive enough to detect differences in both low and high-affinity interactions of HBDs with modified histone peptide tails. We discovered that the hydrophobic clasp (V10, L49) characteristic of the CBX subgroup CBX2, 4, 6, 7, and 8 can be substituted with isoleucine (I) and leucine (L) residues to support H3K27me3 binding. An additional substitution of K33 with glutamic acid (E) that enhanced non-specific binding of CBX7 with H3K27me3^15^ supported stronger H3K27me3-specific binding of the CBX8 clasp variants. Our work has produced variants with up to 5-fold improvement in specific binding to H3K27me3, which is higher than any CBX variant reported to date.

## Results

### Polycomb chromodomains from CBX8 show the most preferential binding of K27me3 over K9me3 compared to other reader domains

In previous work, we have successfully used PCDs to build fusion transcriptional regulators called synthetic reader-actuators (SRAs) that bind H3K27me3 and activate epigenetically-repressed genes in cancer cell lines. The SRA interacts specifically with H3K27me3, and shows no interaction with H3K9me3 *in vitro*^19^. We demonstrated that adding an extra copy of the CBX8 polycomb chromodomain (M1 - R62) with a short glycine-serine linker to a fusion protein increased binding (apparent Kd) by 2-fold in an ELISA-style assay^19^. Binding with H3K27me3 increased, while cross-reactivity with H3K9me3 did not increase^19^. Here, we compared CBX8 PCD to other recently reported engineered histone methyl-lysine-binding proteins derived from CBX7^15^ and CBX1^9^, as well as the BAH domain from BAHCC1^20^ in the same assay (**Fig. 1**).

**Figure 1.**
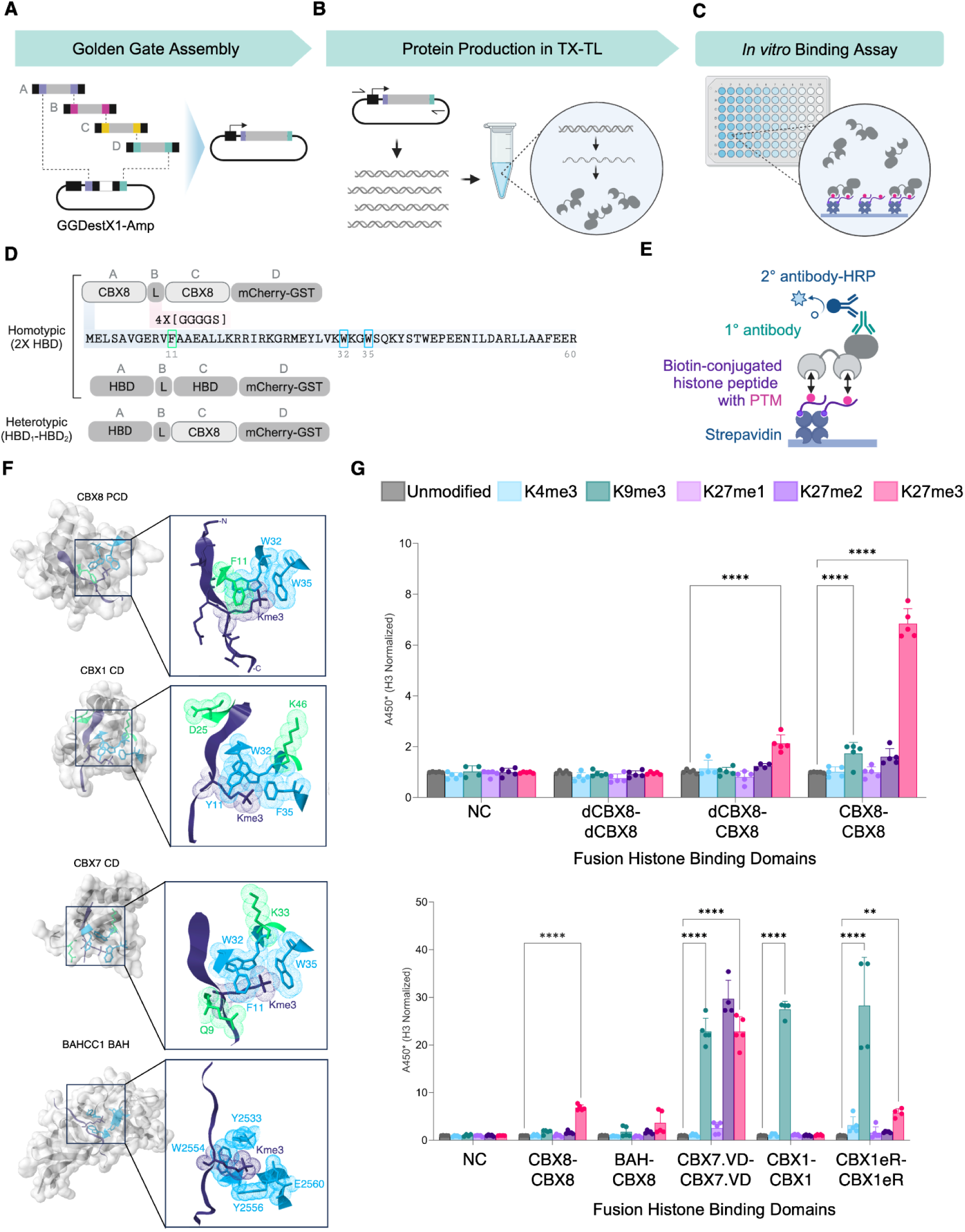
Polycomb chromodomains from CBX8 show the most preferential binding of K27me3 over K9me3 compared to other reader domains. A. Golden Gate assembly is used to build bivalent fusion protein-encoding open reading frames (ORFs) in vector GGDestX1-Amp downstream of promoter OR2-OR1-Pr^18^. B. A gene fragment that includes the promoter and ORF is PCR-amplified and used as a template for cell-free transcription and translation (TX-TL) of the fusion protein. C. The TX-TL product is diluted in blocking buffer, incubated with histone tails, and detected in an ELISA-style assay. D. Schematic of homotypic and heterotypic fusion protein constructs. CBX8 = polycomb chromodomain (PCD) from human CBX8, HBD = CBX8 variant or other histone binding domain, L = 4X [GGGGS] linker. E. Histone binding immunoassay uses streptavidin-immobilized biotin-conjugated histone peptides as target ligands for cell-free expressed fusion proteins. High HRP signal from α-mCherry antibody indicates strong affinity of fusion proteins for histone peptides. F. Structure of human CBX8, CBX7, CBX1 PCD and BAHCC1 BAH domains (RCSB PDB 3I91, 4X3K, 6D07 and 6VIL) with hydrophobic binding pocket residues highlighted in light blue. Residue substitutions in engineered variants (sea green) are as follows: dCBX7 - F11A; CBX7.VD - Q9D, K33E; CBX1eR - K43A, D59F. G. ELISA binding results for wild-type and engineered H3K27me3-binding fusion proteins. Dots = A450 from 5 technical replicate wells, bars = mean of 5 technical replicate wells, error bars = standard deviation, NC = ???.

To determine the degree to which each domain could support specific interaction of a fusion protein with its target histone post translational modification, we built ORFs that encoded a histone binding domain (HBD) at position A, a 4X [GGGGS] linker at position B, another HBD at position C, and an mCherry plus GST tag at position D using our single-pot Golden-Gate assembly method (**Fig. 1A**). Recombinant HBD fusion proteins were expressed in cell-free transcription and translation (TX-TL) reactions^21^ optimized for linear template DNA^22^ and screened for H3K27me3 binding in our Cell-Free Histone-Binding Immunoassay” (CHIA) (**Fig. 1B, C**). We used a bivalent format (2X HBD) because this type of HBD fusion protein is typically used in chromatin probing and pull-down assays ^23–25^. Homotypic proteins include identical HBDs, and heterotypic proteins include two different types of HBDs (**Fig. 1D**). The histone-binding immunoassay detects low affinity histone PTM binding by measuring the amount of fusion protein that remains bound to an immobilized histone peptide (**Fig. 1E**).

The homotypic bivalent fusion that contains wild-type CBX8 PCD showed strong preferential binding to H3K27me3 over partially methylated K27 (me2, me1), K9me3, and K4me3 (**Fig. 1G**). The loss-of-function mutation (F11A) diminished binding in dead-CBX8 (dCBX8) controls, verifying the contribution of the aromatic pocket ^12^ to its interaction with H3K27me3. A 160 a.a. domain from BAHCC1, which contains a 4-residue aromatic cage, was reported to show stronger affinity for H3K27me3 over CBX7 in a recent study^20^. Here, we observed that replacement of a wildtype CBX8 PCD with the BAH domain from BAHCC1 (BAH-PCD) shows specific binding to H3K27me3 with little cross-reactivity to H3K9me3. However, binding was slightly reduced compared to the 2xCBX8 fusion. CBX7.VD was discovered in a phage display screen for strong binders of H3K27me3^15^. This variant includes a substitution near the aromatic cage, where K33 is replaced with glutamic acid (E), and Q9 is replaced with aspartic acid (D) (**Fig. 1F**). A homotypic bivalent fusion containing CBX7.VD, showed strong binding to H3K27me3, and high levels of off-target binding with H3K9me3 and H3K27me2. The cross-reactivity observed in our CHIA assay was also observed for monovalent CBX7.VD in a phage ELISA assay reported by Veggiani et al^15^.

To determine if CHIA could also be used to measure H3K9me3-specific binding, we tested the wildtype CBX1 CD and a recently reported variant called the “eReader” (CBX1eR)^9^. Both wild-type CBX1 and CBX1eR homotypic bivalent fusions exhibited specific binding to H3K9me3. In our CHIA assay, CBX1eR also binds non-specifically to H3K27me3 (**Fig. 1G)**. Replacing the positively charged lysine (K) near the aromatic cage with a negatively charged glutamic acid (E) in CBX7 or aliphatic alanine (A) in CBX1 enhanced the apparent affinity of these proteins for their targets, but also resulted in off-target binding. These results raise the question of whether it is possible to engineer a CBX protein with enhanced affinity for H3K27me3 without introducing cross-reactivity with H3K9me3 or partially methylated forms of Kme27. Therefore, we investigated other potentially tunable features in CBX8 that might contribute to its ability to distinguish K27me3 from K9me3.

### Identification of H3K27me3-Selective Chromodomains in Site-Directed Designed CBX8 Library

To identify novel CBX8 PCD variants with preferential affinity for H3K27me3, we focused on five sites, V10, A12, L49, D50, and R52, which form a histone groove that lies just outside of the conserved hydrophobic pocket (F11, W32, and W35) that interacts with histone H3K27me3 (**Fig. 2A**). A crystal structure study of the Drosophila ortholog Pc suggested that residues at these positions coordinate the interaction of the PCD with the N-terminal tail of histone H3 through hydrogen bonds and van der Waals contacts on the N-terminal side of K27me3^13^. The relative positions of these five residues in Drosophila Pc are conserved in human CBX2, 4, 6, 7, and 8^16^. Aliphatic residues V10 and L49 form a hydrophobic clasp near histone H3A24 (**Fig. 2A**). Replacement of either hydrophobic residue with a polar residue disrupted binding with the H3 ligand. The R-groups of D50 and R52 coordinate hydrogen bonds with the carbonyl groups at histone H3 T20 and L22. Replacement of D50 with cysteine, and R52 or L49 with aspartic acid abrogated binding with H3 tail peptides. These previous studies provide strong evidence for the requirement of contacts of the PCD with histone residues along the N-terminal side of H3K27me3. However these loss-of-function experiments leave open the question of whether amino acid substitutions can maintain or enhance the affinity of PCD for H3K27me3.

**Figure 2.**
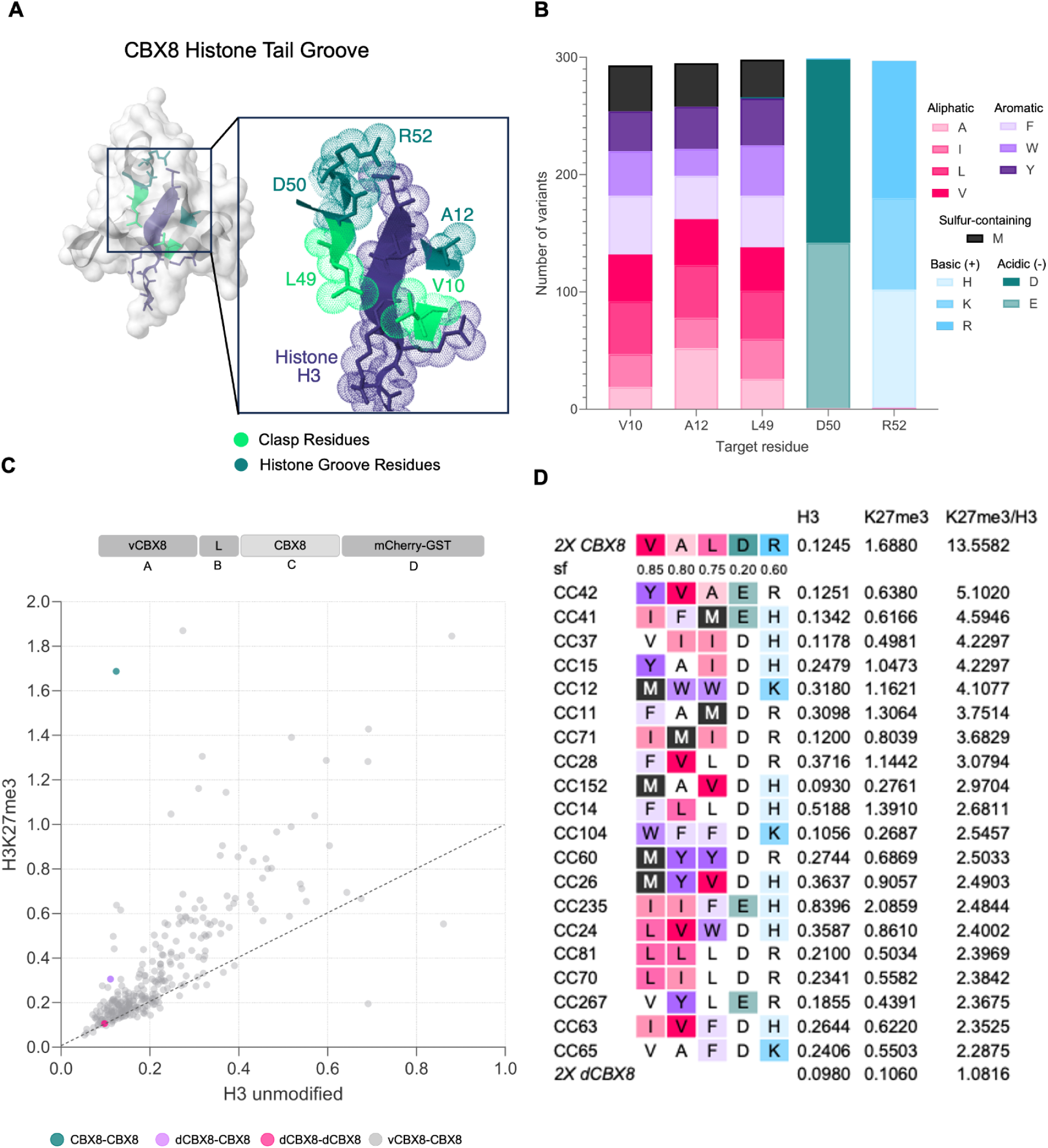
Identification of H3K27me3-Selective Chromodomains in Site-Directed Designed CBX8 Library. A. Structure of human CBX8 PCD domain (RCSB PDB 3I91). Substituted residues in the mutagenesis library are highlighted in teal and green. B. Distribution of residue substitutions in CBX8 library variants. C. Schematic of heterotypic fusion protein ORFs, which encode a variant CBX8 PCD (vCBX8), a 4X [GGGGS] linker, a wild-type CBX8 PCD and a mCherry-GST tag. In the scatter plot, each dot represents the mean A450 from 2 technical replicate ELISA wells. D. Non-specific H3 binding versus on-target H3K27me3 binding signals (mean A450) for 20 CBX8 library variants with the highest on-target binding ratios. Amino acid substitutions compared to wildtype CBX8 are shaded by color as in panel B. Unchanged residues are not shaded. sf = substitution frequency.

We looked for naturally occurring variations by aligning publicly available (Uniprot database) CBX8 ortholog sequences from 62 different species. We observed nearly complete conservation of the aromatic F/Y W W cage and at human positions V10, A12, L49, D50, and R52 (**Supplemental Fig. S1**). Sequences from one cnidarian and two arachnid species contain alternative residues at human positions V10 and L49 (hydrophobic clasp). These include *E. diaphana* (I10 and F49), *A. ventricosus* (F49) and *A. bruennichi* (F49), suggesting tolerance for clasp substitutions as long as the residues are hydrophobic. In these species, human position A12 (hydrophobic) contained charged arginine (R) or threonine (T), which implies greater flexibility in the residues that can occupy this site. The *E. diaphana* sequence includes the polar residue serine (S) at human position R52. Taken together with previous substitution studies, position R52 might require a positively charged or polar residue.

Our evolutionary analysis only provided very limited insights into the diversity of functional CBX8 PCD orthologs. Therefore, we built a library of CBX8 variants with substitutions at these sites. To generate a library of reasonable size for screening, and to avoid amino acid substitutions that might radically alter protein folding and function, we used biochemically conservative substitutions. We replaced V10, A12, and L49 with aliphatic residues (A, F, I, L), aromatic (M, V, W, or Y), or aliphatic sulfur-containing methionine (M). We replaced D50 with a negatively charged acidic residue (E), and replaced R52 with positively charged basic residues (H or K). Quantitative sequencing of the final 3072-variant library of linear fragments confirmed equal representation of all codon substitutions prior to Golden Gate assembly (**Supplemental Fig. S2**).

To ensure adequate library coverage, over 300 CBX8 fusion variants were screened against H3K27me3 using our histone binding ELISA. Amino acid substitutions were relatively evenly dispersed at each position. Aliphatic substitutions accounted for 49% of the residues, while aromatic residues comprised 39% at positions V10, A12, and L49 (**Fig. 2B**). Many variants that demonstrated increased binding to H3K27me3 also exhibited off-target binding to unmodified H3, consistent with previous findings in engineered reader domains, where enhanced affinity often compromises specificity (**Fig. 2C**).

Compared to the homotypic fusion with wildtype CBX8, none of the heterotypic candidates from our screen showed stronger binding to H3K27me3 relative to H3 (**Fig. 2D**). However, many vCBX8-CBX8 fusions maintained some level of affinity for H3K27me3, in contrast to the loss of affinity observed for the heterotypic aromatic cage mutant dCBX8-CBX8. For the variants with the top 20 highest H3K27me3/ H3 binding signal ratios, the most common substitutions appeared at the first three positions V10, A12, and L49, which contain hydrophobic residues. At V10, aliphatic residues (50% V, I, M, or L) appeared at the same frequency as aromatic residues (F, W, Y) with no occurrences of alanine (A). Residues at A12 were predominantly aliphatic (65% A, I, L, or V). To further investigate how substitutions affect binding affinity, we grouped variants based on the occurrence of substitutions at each position (**Supplemental Fig. S3**). Within each group we compared the median H3K27me3 binding values for variants with specific substitutions versus variants with a wildtype residue. For instance, variants that contained an isoleucine (I) substitution at V10 had a slightly higher median value than variants with no change at V10. Overall, median increases were only observed for sub-groups with substitutions at the first three hydrophobic positions, V10 (V10I), A12 (A12V), and L49 (L49A, L49I), whereas slight decreases were observed for groups with substitutions at D50 and R52. The results from our library screen suggest that substitutions at the hydrophobic clasp region are tolerated by the CBX8 PCD, while substitutions at the charged residues D50 and R52 may compromise binding. Next, we generated and tested a new set of variants that only carried specific substitutions in the hydrophobic clasp.

### Discovery of novel amino acid substitutions in CBX8 that support H3K27me3 binding

The clasp motif within CBX proteins has been studied as a potential contributor to CBX selectivity^16^ with a marked difference between CBX proteins that bind K27me3 versus K9me3. H3K27me3-binding CBX family members (CBX2,4,6,7 and 8) contain hydrophobic, non-polar residues, valine (V) and leucine (L), in the clasp while H3K9me3-binding CBX family members (CBX1, 3, 5, and 9) contain polar, negatively charged residues, glutamate (E) and aspartate (D), in the clasp. This pattern is also observed in non-CBX H3K9me3-binding and H3K27me3-binding chromodomains^26,27^ (**Fig. 3A**). Although the surrounding regions of K9 and K27 are similar, both featuring an ARKS motif, the residues flanking this motif differ. K27 is flanked by non-polar and uncharged alanine residues (A24, A29) whereas K9 is flanked by polar, uncharged threonine residues (T6 and T11) (**Fig. 3B**). We hypothesized that polar clasp residues in H3K9 binders strengthen hydrogen bonding with adjacent threonine residues thereby enhancing selectivity for K9. Based on this reasoning, we hypothesized further increasing the hydrophobicity of the clasp in CBX8 proteins could potentially enhance their selectivity for K27 over K9.

**Figure 3.**
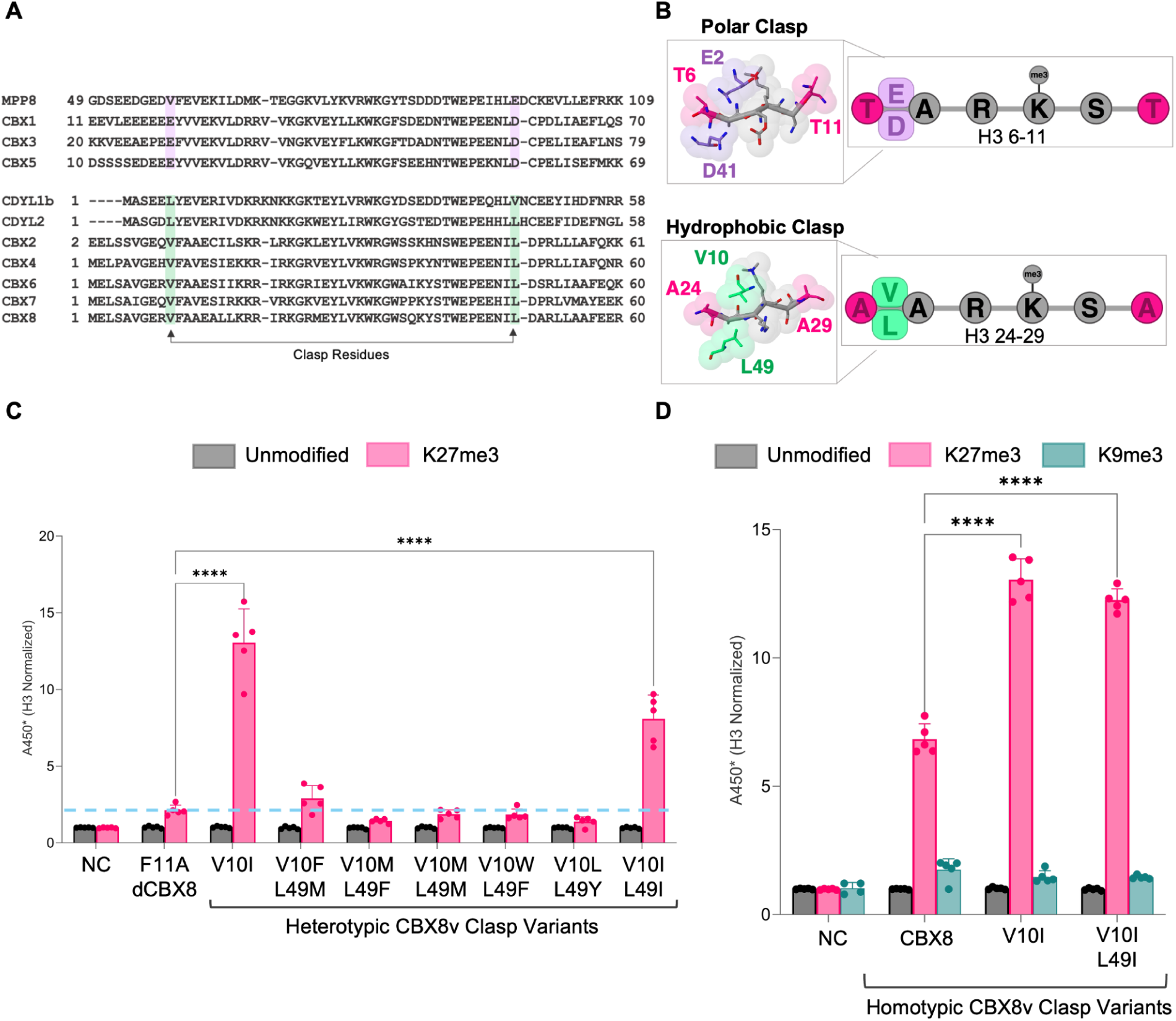
Discovery of novel amino acid substitutions in CBX8 that support H3K27me3 binding. A. Sequence alignment of select chromodomains. H3K9me3-binding domain clasp residues are highlighted in light purple and H3K27me3-binding domain clasp residues are highlighted in sea green. B. Structural diagram of polar clasp interaction with AARK9S motif (top) and hydrophobic clasp interaction with AARK27S motif (bottom) within histone H3 tail. Polar clasp residues (E2 and D41) are highlighted in light purple and hydrophobic clasp residues (V10 and L49) are highlighted in sea green. Pink highlighted residues denote start and end of each A-R-K-S motif on histone H3 tail. C. Replicate ELISA binding data for heterotypic CBX8 clasp variants. The blue dotted line marks the mean value for heterotypic dCBX8/ H3K27me3 binding. (**** *p* ≤ 0.0001) D. Replicate ELISA binding data for select homotypic CBX8 clasp variants. For panels C and D, dots = A450 from 5 technical replicate wells, bars = mean of 5 technical replicate wells, error bars = standard deviation.

To explore the idea that hydrophobic residues contribute to a H3K27me3-compatible clasp, we closely inspected variants from our library that contained hydrophobic substitutions (L,V, F, I) at one or both clasp positions V10 and L49. Both phenylalanine (100) and isoleucine (99) had a higher hydrophobicity index than leucine (97) and valine (76) at pH 7^28^. The potential functionality of these substitutions is also supported by naturally occurring CBX8 clasp variations we identified in one cnidarian (I10, F49) and two arachnids (F49) (**Supplemental Fig. S1**). Tryptophan was excluded due to its bulky aromatic side chain, which could disrupt the clasp motif. As previously reported, we observed mutation to clasp residues is context dependent, and can significantly alter binding affinity^16^. For instance, mutation to V10F and L49M improved binding 1.4-fold over heterotypic dCBX8. However, swapping these positions to V10M and L49F abrogated binding to levels comparable to homotypic dCBX8 (**Fig. 3C**). Conversely, mutation V10I improved binding by 6.1-fold, and both V10I and L49I mutations resulted in a 3.8-fold increase in binding affinity compared to the dCBX8 fusion. These findings suggest V10I mutation is the major contributor to enhanced binding affinity.

We previously showed that CBX8 PCD binding is cooperative, with bivalency increasing avidity in a non-linear fashion^19^. To account for potential cooperativity of homotypic readers, we generated and tested homotypic CBX8 V10I and CBX8 V10I L49I fusion proteins in CHIA. Both homotypic CBX8 V10I and CBX8 V10I L49I exhibited increased H3K27me3 binding compared to homotypic wild-type CBX8 (**Fig. 3D**). To assess whether these mutations affected specificity for H3K27me3, we tested the variants against H3K9me3. Both variants had the same amount of binding for H3K9me3 as wild-type CBX8 (**Fig. 3D**). Taken together, these results suggest that the clasp variants maintained specificity. We next investigated whether substitutions closer to the binding pocket could further improve binding affinity.

### A substitution at K33 enhances binding of CBX8 to H3K27me3 without cross-reactivity with H3K9me3

Previous studies of polycomb chromodomain structure and function suggest that K33 in human CBX6, 7, and 8, which appears near the aromatic cage that surrounds the trimethylated histone lysine, contributes to histone binding affinity. In synthetic variant CBX7.VD^15^ K33 is replaced with glutamic acid (E) (**Fig. 1A**). The BAH domain contains glutamic acid (E2560) within the aromatic cage reported by Fan et al^20^, and showed enhanced affinity for H3K27me3 compared to wild type CBX7. In the previously reported phage display screen that identified CBX7.VD^15^, a candidate derived from CBX8 contained substitutions K33E and Q9D showed increased affinity for H3K27me3, but also bound tightly with H3K9me3 in a phage ELISA assay. These observations prompted us to investigate whether the K33E substitution in wildtype CBX8 and the clasp variants could enhance binding without compromising specificity.

To determine the effect of K33E in the context of wildtype CBX8 PCD, we used CHIA to test a homotypic bivalent fusion protein, 2X CBX8 K33E, that carried this substitution. This variant showed significantly increased binding with H3K27me3 compared to the wildtype control 2X CBX8, as well as 2X CBX7.VD (*p* < 0.001) (**Fig. 4A**). Off-target binding with H3K9me3 was substantially reduced (∼5.8-fold) for 2X CBX8 K33E compared to 2X CBX7.VD. Introduction of K33E into clasp variant V10I resulted in H3K27me3 binding that surpassed wildtype CBX8 and CBX8 K33E. Off-target binding with H3K9me3 also increased, but this variant maintained a strong preference for H3K27me3. A fusion protein with clasp substitution V10I L49I and K33E showed reduced H3K27me3 binding compared to the variant V10I K33E, but the binding signal was stronger than that for wildtype CBX8 and off-target binding for H3K9me3 was not significantly increased (*p* = 0.7046).

**Figure 4.**
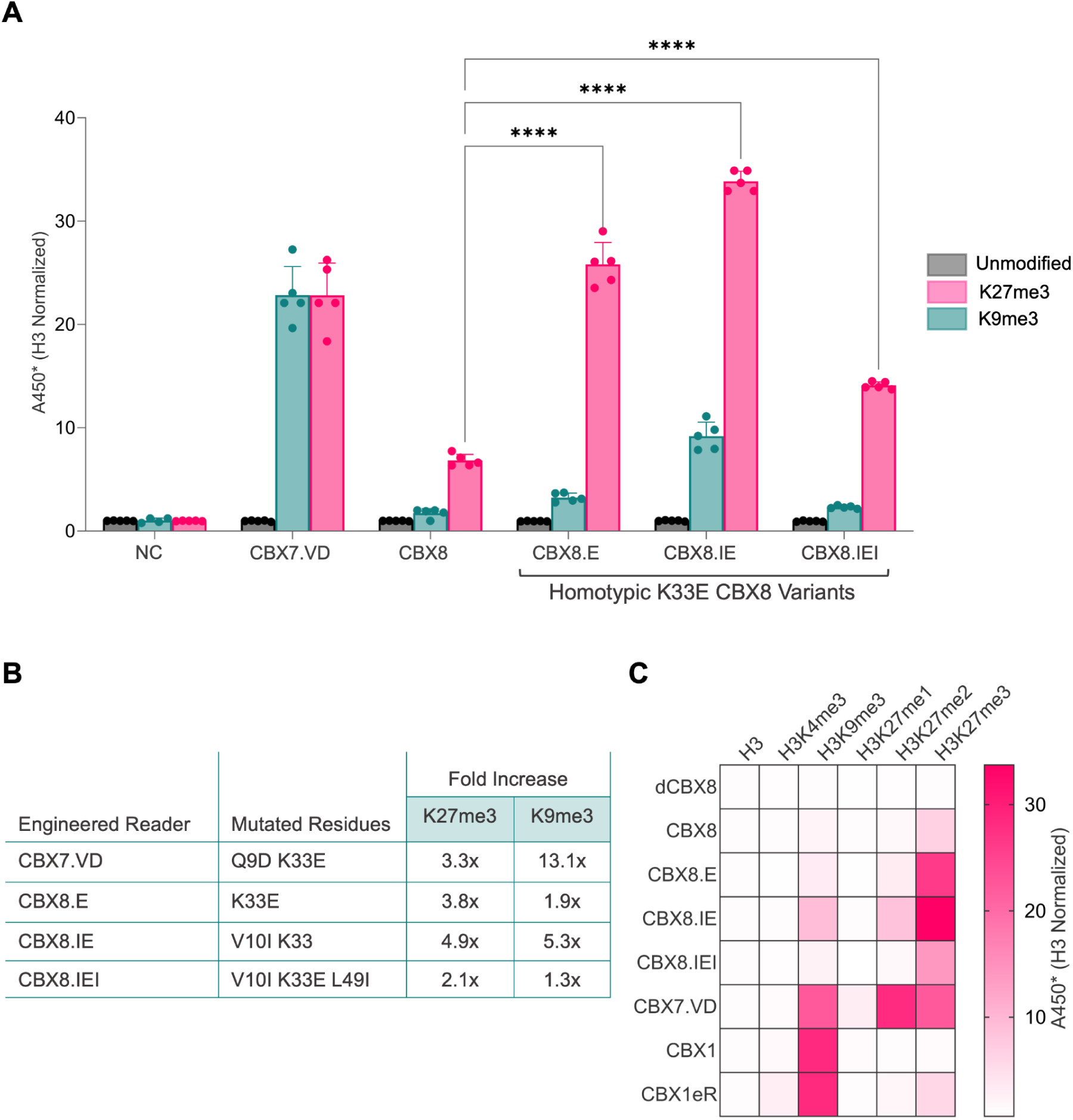
A substitution at K33 enhances binding of CBX8 to H3K27me3 without cross-reactivity with H3K9me3. A. ELISA binding data for top-performing homotypic CBX8 variants after the introduction of K33E. Dots = A450 from 5 technical replicate wells, bars = mean of 5 technical replicate wells, error bars = standard deviation. (**** *p* ≤ 0.0001) B. Specificity profiling of homotypic CBX fusion proteins against a panel of unmethylated and methylated histone peptides. Absorbance signal for each histone PTM is normalized to the average signal from the H3 unmodified (negative control) ELISA wells. C. Fold increase of binding to H3K27me3 and H3K9me3 for top-performing CBX8 variants compared to wild-type CBX8. Values for H3K27me3-reader CBX7.VD and H3K9me3-readers CBX1 and CBX1eR are included for comparison.

A summary of the overall performance of our novel CBX8 variants compared to other methyl-lysine-binding engineered proteins is presented in **Fig. 4B and 4C**. The most common off-target binding for all proteins (>3.7 H3-normalized signal) occurred for H3K9me3 (**Fig. 4B**). Specific interaction of a CBX1 fusion with K9me3 and not K27me3 supported the validity of CHIA for the detection of protein interactions with K9me3. We observed little to no binding with H3K4me3, another biologically important lysine methylation mark. For undermethylated K27 peptides (K27me1 and K27me2) we observed minimal binding (0.9 - 8.2 H3 normalized signal) by the CBX8 variants K33E, V10I K33E, and V10I K33E L49I, whereas CBX7.VD binding to H3K27me2 surpassed its H3K27me3 binding signal (**Supplemental Fig. S4**). Serial dilution experiments confirmed that the relative performance of the engineered variants was maintained within the linear range of the CHIA assay (**Supplemental Fig. S5**). Among the engineered variants, 2X V10I K33E L49I (CBX8.IEI) combined the strongest H3K27me3 binding with minimal recognition of H3K27me1/2, while also exhibiting one of the highest H3K27me3:H3K9me3 specificity ratios. Based on this favorable balance of affinity and selectivity, we prioritized CBX8.IEI for subsequent cell-based characterization.

### An engineered CBX8 variant fused to GFP shows nuclear punctae in cultured human cells

To investigate the specificity of engineered CBX-derived histone-binding domains for H3K27me3 in the context of whole chromatin in live cells, we built a set of enhanced green fluorescent protein (eGFP) fusions that carried N-terminal tandem homotypic domains. We expected these to function as fluorescent probes^5,6,15,23–25^, where bright punctae indicate areas of H3K27me3 enrichment, a pattern seen for immunostaining of H3K27me3^29,30^ and PRC-enriched Polycomb bodies^31,32^. We placed the fusion proteins under the control of a doxycycline-inducible TetTA-regulated promoter in the vector pSBtetTA-luc2GG (**Fig. 5A**) and generated stable transgenic cell lines, including triple negative breast cancer (TNBC) BT-549 which are associated with high levels of H3K27me3 generated by hyperactive histone methyltransferase EZH2^33–38^. Others including HCC1806 (TNBC), HEK293, and Jurkat were included to assess generalizability of the results. After two days of activation of the transgene with doxycycline, probes carrying histone domain CBX8.IEI showed bright punctae over a dimmer background in BT-549, HEK293, and Jurkat, whereas the signal was more diffuse in HCC1806 (**Fig. 5B, Supplemental Fig. S6**). The negative control dCBX8, which has a mutated H3K27me3 binding pocket, showed a mostly diffuse signal throughout the nucleus. We also observed a few large bright areas in these nuclei, presumably representing nucleolar accumulation of the fusion protein, a behavior previously reported for GFP fusions containing the same SV40 nuclear localization signal^39^. *[Describe immunofluorescence staining results]* (**Fig. 5D**). Overall, these results demonstrate that the H3K27me3-binding PCD is required for the formation of distinct nuclear puncta, whose morphology is similar to the inactive X, and H3K27me3-enriched nuclear foci reported previously by immunofluorescence^40–42^. Next, we sought to experimentally determine if CBX8.IEI-GFP relies on the active generation of H3K27me3 by EZH2.

**Figure 5.**
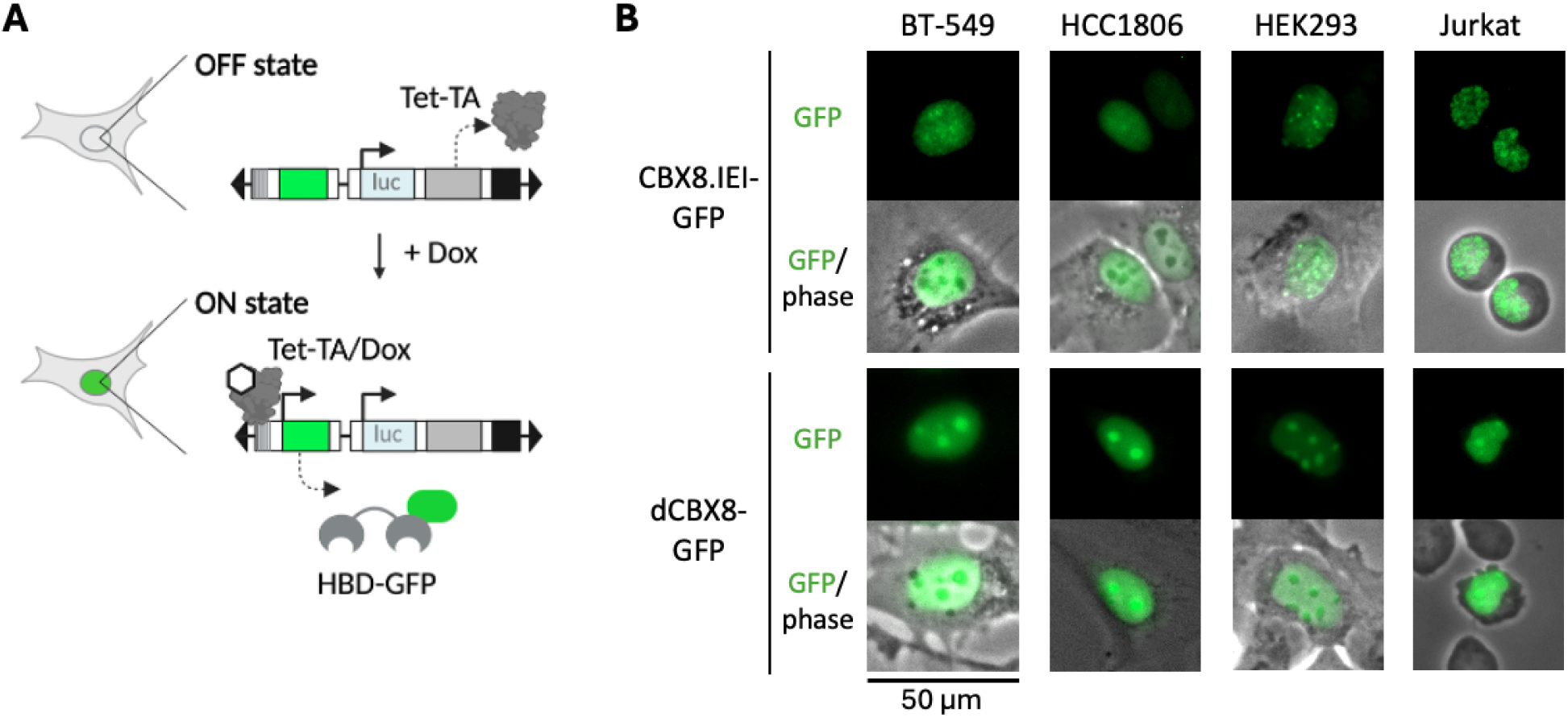
CBX8.IEI-GFP shows nuclear punctae in cultured human cell lines. A. Vector pSBtetTA-luc2GG carries a constitutive RPL13a promoter that drives firefly luciferase, Tet-3xVP16 activator (TetTA), and puromycin resistance expression, and the expression of fusions containing HBDs, green fluorescent protein, and nuclear localization sequence (HBD-GFP) is driven by doxycycline-bound TetTA. B. Representative images of CBX.IEI-GFP probe localization in BT-549, HCC1806, HEK293, and Jurkat cells, compared with non-binding dCBX8-GFP proteins.

### Modulation of EZH2 activity abrogates engineered CBX8 binding in breast cancer cells

We hypothesized that the inhibition of EZH2 would abrogate punctate signals from on-target binding to H3K27me3 and/or H3K27me2^43^, while any off-target signals associated with binding to H3K9me3 and other marks would remain unchanged (**Fig. 6A**). Similar to probes with CBX8.IEI, probes carrying histone domains CBX7.VD and CBX8 (w.t.) showed bright punctate signals over a dimmer background after two days of activation of the transgene with doxycycline (**Fig. 6B**). We scored probe signal patterns using manual qualitative assessment of 60 nuclei per condition, computational image segmentation of nuclear punctae (dots), and a computational classifier for rapid, quantitative scoring of hundreds of nuclei. Nuclei were classified as “compartment,” showing large areas of unbound probe typical of dCBX8-GFP, or “speckled,” showing many smaller bright foci. Following EZH2 inhibition, CBX8.IEI probes exhibited a more complete loss of the speckled phenotype than wild-type CBX8 (**Fig. 6C**), consistent with stronger dependence on H3K27me3 for chromatin localization. In contrast, a substantial fraction of nuclei expressing wild-type CBX8 retained speckled localization after EZH2 inhibition. In contrast, CBX7.VD punctae appeared to be insensitive to EZH2 inhibition, suggesting substantial off-target, non-H3K27me3 binding. This idea is consistent with our *in vitro* data, where CBX7.VD showed strong binding with H3K9me9 (**Fig. 4**).

**Figure 6.**
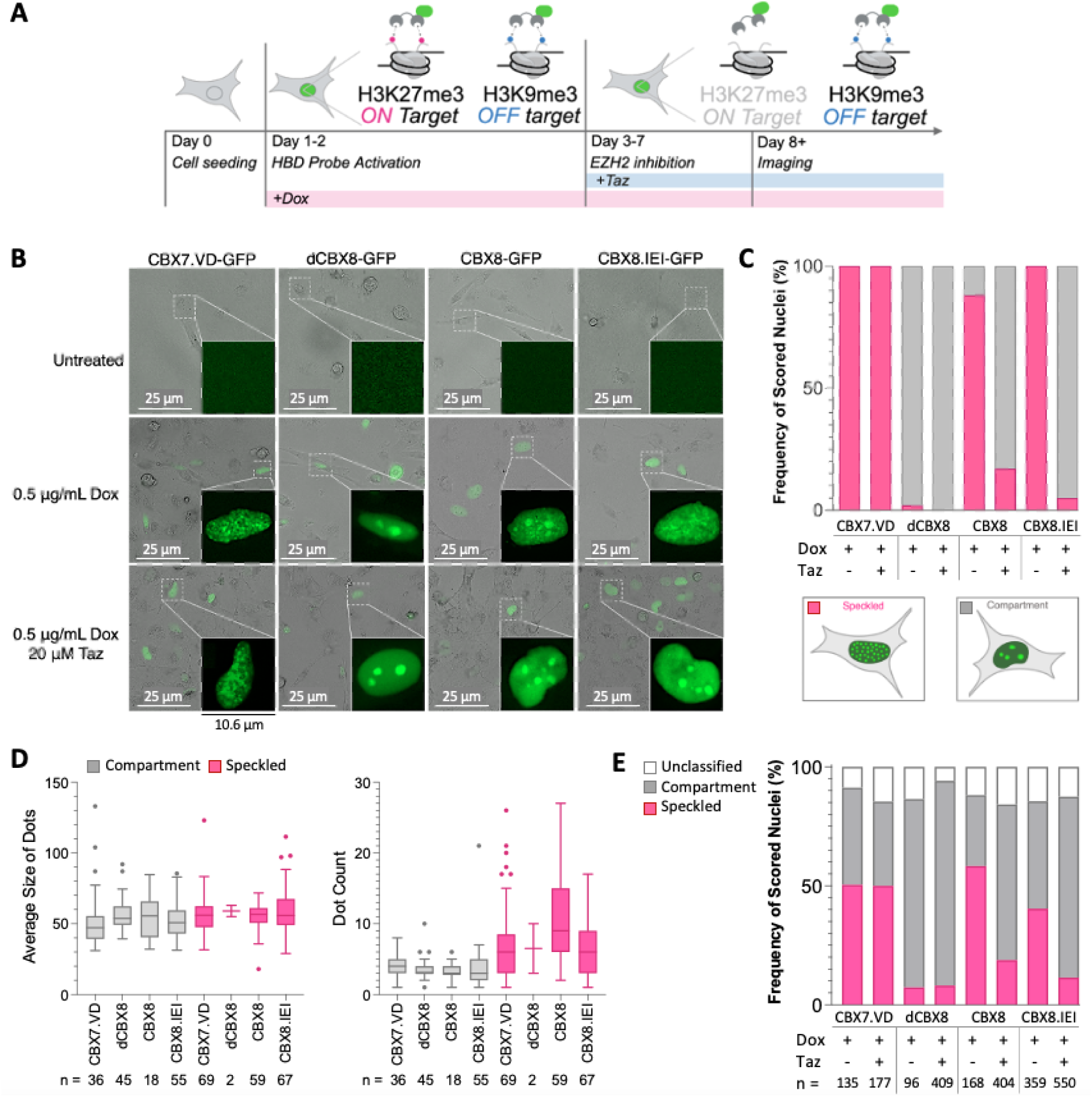
Modulation of EZH2 activity abrogates engineered CBX8 binding in breast cancer cells. A. Experimental design. BT-549 cells expressing HBD-GFP probes were seeded (Day 0), induced with 0.5 µg/mL doxycycline (Dox, Days 1-2), treated with the EZH2 inhibitor tazemetostat (Taz, 20 µM; Days 3-7), and imaged on Day 8. B. Fluorescence live imaging of HBD-GFP expression in untreated (top), 0.5 µg/mL doxycycline-treated (middle), and 0.5 µg/mL Dox + 20 µM tazemetostat-treated cells (bottom). Scale bar, 25 µm. C. Manual scoring of fluorescence signal patterns (left) in CBX7.VD, dCBX8, CBX8, CBX8.IEI nuclei (*n* = 60 per condition) before and after tazemetostat treatment (left). D. Comparison of GFP “dots” identified by Cellpose in nuclei classified by the machine learning model as Compartment or Speckled (confidence = 1.0). n = number of nuclei in each box plot group. E. Computational classifier results are shown for nuclei with confidence scores of 0.59 - 1.0.

We used computer-aided image segmentation to characterize GFP signal patterns across larger cell populations. Compartment and speckled nuclei exhibited distinct distributions of dot counts per nucleus, whereas differences in average size (area) of dots per nucleus were less pronounced (**Fig. 6D**). This likely reflects the manner in which Cellpose identified GFP-positive regions, with segmentation boundaries influenced by signal intensity rather than the broader spatial organization of nuclear GFP patterns (**Supplemental Fig. S7A**). The machine-learning classifier remained broadly concordant with manual scoring (**Supplemental Fig. S7B**, **Appendix A3**), indicating that phenotype classification was driven primarily by differences in signal organization and dot abundance rather than dot size alone.

### Tumor microenvironment-related cues modulate CBX8.IEI-GFP signal patterns

Next, we sought to determine whether tumor microenvironment-associated cues modulate probe localization independently of pharmacologic EZH2 inhibition. Specifically, we examined the effects of extracellular matrix stiffness and transforming growth factor (TGF)-β1 stimulation on CBX8.IEI-GFP signal patterns in BT-549 cells cultured on fibronectin-coated hydrogel substrates. This experiment was motivated by our recent work demonstrating that, under conditions of increased matrix stiffness and TGFβ1 activation, EZH2 exhibits enhanced nuclear localization and activity^44^. Qualitative comparison of cells cultured on soft or stiff matrices revealed modulation of the GFP signal pattern in response to TGFβ1 treatment (**Fig. 7A**). Cells on soft matrices showed a diffuse GFP signal with unclear punctae, but this pattern was distinguishable from the non-binding control construct (dCBX8-GFP). TGFβ1 treatment showed no effect for soft-matrix samples. Stiff matrix samples showed heterogeneous patterns with more intense GFP areas against a diffuse background, and TGFβ1 treatment intensified the number and brightness of these areas. Similarly, cells grown on fibronectin-coated glass showed intensified GFP signals with TGFβ1 treatment.

**Figure 7.**
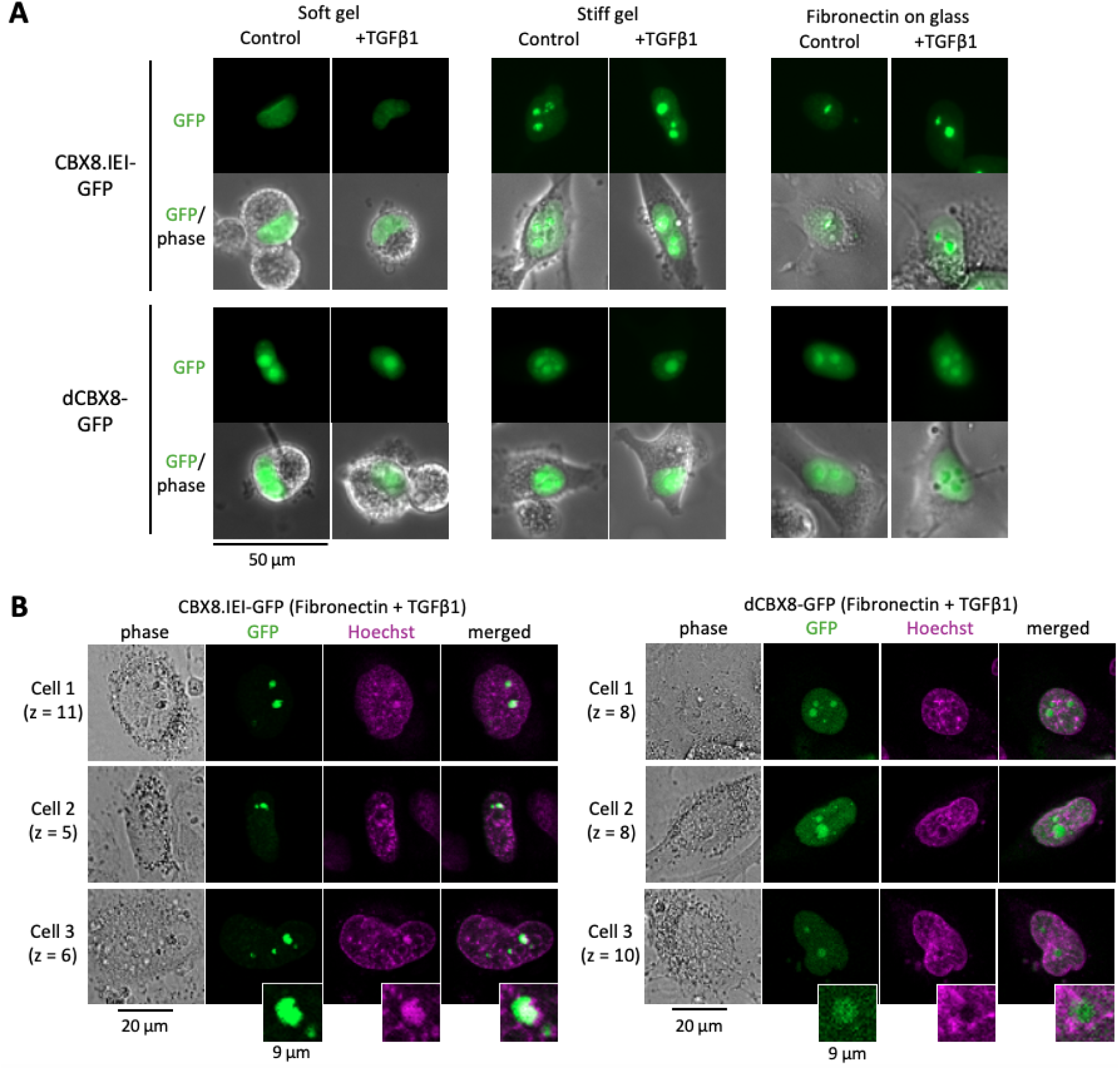
Tumor microenvironment-related cues modulate CBX8.IEI-GFP signal patterns. A. Representative fluorescence images of live BT-549 cells expressing CBX8.IEI-GFP or the non-binding control probe (dCBX8-GFP) cultured under defined extracellular microenvironment conditions. Cells were grown on soft hydrogel matrices, stiff hydrogel matrices, or fibronectin-coated glass, with or without TGFβ1 stimulation. Merged GFP/phase images are shown below the corresponding fluorescence images. B. Representative single-plane confocal images of fusion protein-expressing BT-549 cells cultured on fibronectin in the presence of TGFβ1, followed by staining with Hoechst 33342. Insets highlight representative bright GFP areas and surrounding nuclear context.

The large, bright GFP regions observed under fibronectin and TGFβ1 conditions were distinct from the smaller puncta seen in cells grown on tissue culture plastic (**Fig. 5, 6**), prompting us to determine whether these structures represented intranuclear, chromatin-associated features rather than probe aggregates. To address this, we examined z-plane series of CBX8.IEI-GFP-expressing cells cultured on fibronectin-coated glass in the presence or absence of TGFβ1. Across z-slices, the intense GFP puncta remained spatially continuous with the surrounding diffuse nuclear GFP signal, indicating that these structures reside within the nuclear volume (**Supplemental Fig. S8**). To further assess their relationship to chromatin, we performed confocal imaging with Hoescht 33342 staining. Bright CBX8.IEI-GFP regions were consistently observed within DAPI-positive nuclear regions, arguing against the interpretation that these structures represent DNA-excluded aggregates (**Fig. 7B**). In contrast, the non-binding control probe (dCBX8-GFP) exhibited a more diffuse distribution that was not preferentially enriched in DAPI-dense regions and was frequently observed in DAPI-dim compartments, consistent with accumulation in chromatin-poor nuclear subregions. Together, these observations support that the large GFP puncta reflect chromatin-associated probe localization and demonstrate that tumor microenvironment-relevant cues modulate CBX8.IEI nuclear organization in an orthogonal, non-pharmacologic context.

## Discussion

We demonstrated that CHIA is an effective system for comparing the strength and specificity of low-affinity wildtype and engineered histone reader domains, specifically those that recognize different methylated lysines on the histone H3 tail. The modularity of Golden Gate cloning allows “single pot” digestion-ligation assembly of fusion proteins that contain different combinations of histone binding domains, potentially those that recognize other histone PTMs (e.g. acetylation, phosphorylation, etc.) that were not included in this study. CHIA could also be easily customized to interrogate protein interactions with other biotin-conjugated ligands, such as single and combinatorial histone PTMs, mixtures of different histone PTMs in a single well, whole nucleosomes, or *in vitro*-methylated DNA. By using a gain-of-affinity approach based on a highly specific but low-affinity histone binding protein, we identified novel engineered CBX8-derived H3K27me3 readers that outperform other engineered variants in affinity and specificity.

We also gained important insights into the structure and function of the H3K27me3-specific polycomb chromodomain (PCD). Although the aromatic cage that recognizes methylated lysine has been the defining feature of the PCD for more than two decades^17^, structural and biochemical studies have established that histone recognition depends on additional functional surfaces, including a histone tail-binding groove^13,16^, an arginine-rich patch proposed to stabilize interactions with the nucleosome core^14^, and a conserved hydrophobic clasp^16^. Our results expand this framework by demonstrating that the hydrophobic clasp is not merely a structurally conserved element but is itself amenable to rational engineering. Specifically, the evolutionarily conserved V10-L49 clasp can be replaced by I-L or I-I combinations while maintaining specificity for H3K27me3 over H3K9me3. In addition, substitution of lysine 33 with glutamic acid adjacent to the conserved aromatic cage strengthens binding in CBX8, consistent with previous observations that analogous substitutions enhance affinity in other chromodomains. Together, these findings suggest that the PCD functions as a compact, multifunctional recognition module whose individual structural elements can be engineered independently to improve histone mark recognition while preserving specificity *in vitro* and in live cells.

## MATERIALS AND METHODS

### Golden Gate assembly of fusion protein genes

A histone binding domain (HBD), linker domain, a second HBD, and GST-tagged mCherry fluorescent protein for positions A, B, C, and D, respectively, were flanked by Golden Gate adaptors as previously described^18^ using the following Q5 HiFi PCR reaction: 5X Q5 reaction buffer (5 μL); 10 mM dNTPs (0.5 μL); 10 μM forward primer (1.25 μL); 10 μM reverse primer (1.25 μL); Q5 HiFi DNA polymerase (0.5 μL); DNA template (0.5 μL); molecular biology grade H_2_O (16.25 μL). Human CBX8 was cloned from our previously reported construct^19^. Plasmid templates from Addgene included CBX1 (ID 180212) and CBX1eR (ID 135089). CBX7.VD and K33E CBX8 templates from Fig. 1 and 5 were synthesized by Integrated DNA Technologies (IDT gBlocks). Primer binding regions are provided in **Supplemental Table S1** and template sequences are provided in **Supplemental Appendix A1**. The products (25 μL) were blunt-end ligated into vector pB2K (PCR-blunt II-TOPO, Thermo #45-024-5) to generate donor plasmids with Kanamycin resistance. Donors were validated by Sanger sequencing (Azenta/ Genewiz) with primer M13R 5’-caggaaacagctatgac. 40 fmol of each donor plasmid and the GGDestX1-Amp destination vector (Addgene #157652) were used for single-pot assembly with BbsI (BpiI) restriction enzyme (Thermo #FERFD1014) and T4 ligase (Quick ligase, NEB #M2200) as previously described^18^. Each ligation product was transformed into *E. coli* DH5α (NEB Turbo, NEB #C2984H) and purified from a 3 mL overnight liquid culture of a single colony (Zymo #D4016).

### Cell-free expression of fusion proteins

Linear PCR products were generated using primers

5’-TTGACATGGTGAAGtCTATCGCACCATCAG and 5’-CGACGATAGTCATGCCCCGCGC, purified using GenElute™ PCR Clean-Up Kit (Sigma-Aldrich #NA1020-1KT) and added to reaction at 390 ng. Cell-free lysates were prepared by the M. Styczynski lab using BL21 DE3 STAR Δlacizya strain of *E. coli* as described previously^21^. Cell-free reactions were set up with the following components: 12 mM Mg-glutamate, 10 mM ammonium glutamate, 130 mM K-glutamate,1.2 mM ATP, 0.85 mM GTP, CTP and UTP, 0.068 mM folinic acid, 0.17 mg/mL tRNA, 0.33 mM NAD, 0.26 mM coenzyme A, 4 mM oxalic acid, 1 mM putrescine, 1.5 mM spermidine, 57 mM HEPES, 2 mM of each of the 20 amino acids, 0.3 M PEP and 27% v/v of cell extract. Chi6 dsDNA was generated by annealing ssOligos 5’-TCACTTCACTGCTGGTGGCCACTGCTGGTGGCCACTGCTGGTGGCCACTGCTGGTGGCCACTGCTGGTGGCCACTGCTGGTGGCCA and 5’-TGGCCACCAGCAGTGGCCACCAGCAGTGGCCACCAGCAGTGGCCACCAGCAGTGGCCACCAGCAGTGGCCACCAGCAGTGAAGTGA at 95°C for 3 mins, chilled to 4°C and used at 2.5 nM in cell-free reactions^22^. All cell-free reactions were carried out in a volume of 10 μL in a 96-well plate (Thermo Scientific #4483348). Fusion protein expression was measured via mCherry fluorescence96x 10 minutes) (533−610 nm) in a QuantStudio 6 thermal cycler set at 29°C (with a signal read after each cycle (16 hrs total).

### ELISA-style binding assays

ELISA-style protein-histone interaction assays were performed as previously described^45^ with modifications. All steps were carried out at room temperature and all incubations and washes were agitated at 800 rpm on an VWR Orbital Shaker 1000. Briefly, clear bottom plates (Greiner bio-one #655101) were coated in 20 ng/μL neutravidin in PBS pH 8.0 overnight at 4 °C. Following 3 washes with 0.2% PBST, plates were blocked with 5% BSA in 0.2% PBST for 30 mins. Plates were then incubated with 1 μM biotinylated histone tails for 1 hr. Histone tails used in this study included H3 A21− G44 (Anaspec #AS-64440-025), H3K27me3 A21−G44 (Anaspec #AS-64367-025), H3K27me2 A21−G44 (Anaspec #AS-64366-025), H3K27me1 A21−G44 (Anaspec #AS-64365-025), H3K9me3 A1-A21 (Anaspec #AS-64360-025), and H3K4me3 A1-A21 (Anaspec #AS-64357-025). Following 3 washes with 0.2% PBST, plates were blocked with 5% skim milk in 0.2% PBST for 30 mins. Cell-free reactions diluted 1:25 in 5% skim milk in 0.2% PBST were incubated in plates for 1 hr. Following washing with 5% skim milk in 0.2% PBST, 1:1000 polyclonal anti-mCherry (Novus Biologicals #NBP2−25158) in 5% skim milk in 0.2% PBST were incubated for 1 hr at room temperature. After addition of 1:1000 rabbit anti-chicken−HRP (RCYHRP Genetel 0.5 mg/mL) in 5% skim milk in 0.2% PBST, plates were incubated for 30 min. Following washing with 0.2% PBST, plates were incubated with 1-step Ultra TMB-ELISA (Thermo-Fisher #34029) for 15 min while protected from light. Reactions were stopped with 2.0 M sulfuric acid, incubated for 2 min, and read at 450 nm on SpectraMax iD3 plate reader. For each sample, the A450 value for each well (four replicates) was divided by the mean A450 value of four H3 (unmodified histone) control wells.

### CBX8 PCD variant library

CBX8 PCD variants flanked by Golden Gate adaptors for position A were generated by Genscript as a combinatorial library with codon substitutions at V10, A12, L49, D50, and R52. Deep sequencing was performed by Genscript to verify that codon representation was evenly distributed (see **Supplemental Fig. S2**). Golden Gate assembly was done using the A-part library (40 fmol linear fragments) instead of plasmid donor A as described for “Golden Gate assembly of fusion protein genes.” Transformation, plating for single colonies, colony PCR with primers GGDestX1-Amp forward 5′-GTGATAATGGTTGCAGCTAGC and GGDestX1-Amp reverse 5′-GGCTTTGCTCGAGTTAGC, plasmid purification, and Sanger sequencing were done as previously described^18^.

### Mammalian expression plasmid construction

The pSBtetTA-luc2GG Golden Gate destination and expression vector (Benchling, https://benchling.com/s/seq-nwX64g96g87pi2WUny44?m=slm-eyZ7vNHXNeiXc0GhNY0b) was built from pSBtetTA-YP^46^. Gibson Assembly (NEB #E2621S) was used to replace YFP with luciferase by ligating PCR amplicons generated from pSBtetTA-YP excluding the YFP ORF (forward 5’-agggcggcaagatcgccgtgggcagtggagctactaacttcag, reverse 5’-atgtttttggcatcttccatggtggcctcaggtgc), and a luciferase ORF from pSBtetGP (Addgene, #60495)^47^ (forward 5’-cctctgcacctgaggccaccatggaagatgccaaaaacattaagaagggc, reverse 5’-aagttagtagctccactgcccacggcgatcttgccgccct). The multi-cloning site (MCS) downstream of TetO-CMV was modified to include a 2X BbsI Golden Gate drop-in site (underlined) by inserting annealed oligos (top 5’-ggccgcatctaga<u>cccgccgccaccatggag</u>gggtcttcgagaagacctactagtagc, bottom 5’-ggccgctactagtaggtcttctcgaagaccc<u>ctccatggtggcggcggg</u>tctagatgc) with sticky overhangs into a NotI-linearized plasmid. Other BbsI sequences were removed using site directed mutagenesis (NEB, #E0554S) with specific primers for luciferase (forward 5’-cccactcgaagatgggaccgccg, reverse 5’-tagaatggcgctgggccc) and BGH polyA (forward 5’-aggattgggatgacaatagcag, reverse 5’-cccccttgctgtcct). Golden Gate assembly^18^ was used to build fusions with parts A, B, and C as before, but with eGFP-NLS-stop at the D position (**Supplemental Table S1**, **Supplemental Appendix A1**). Plasmid sequences were verified with Sanger sequencing (Azenta Genewiz) and nanopore sequencing (Plasmidsaurus).

### Cell Culture and Transgenic Cell Line Generation

Cells were cultured in complete media supplemented with 10% (v/v) tetracycline system approved fetal bovine serum (Thermo #A4736401) and 1% penicillin-streptomycin (Thermo #15140-122) using base media and supplements as follows: BT-549 (ATCC #HTB122) in RPMI-1640 (Corning #10-104-CV) and 0.8 µg/mL insulin (Thermo #12585014); HCC1806 (ATCC #CRL-2335) in RPMI-1640 and 2 mM L-glutamine (Thermo #25030081); HEK293 (ATCC #CRL-1573) in DMEM high glucose (Thermo #11965092); and Jurkat (ATCC #TIB-152) in RPMI-1640 and 0.1 mM non-essential amino acids (Thermo Fisher 11140-050). Cells were passaged using standard techniques; adherent cells were washed with 1X DPBS without calcium and magnesium (Corning #20-031-CV) and detached using 0.25% trypsin-EDTA (Thermo #25200056). Prior to transfection, 2×10^5^ cells were seeded in 0.5 mL penicillin-streptomycin-free culture medium in a 12-well microplate. Transfections were performed by adding DNA-Lipofectamine complexes (260 µL) dropwise to each well: 1.6 µg plasmid DNA, 0.1 µg helper SB100X plasmid DNA, 2.5 µL Lipofectamine LTX (Thermo #15338030), and 1.3 µL PLUS Reagent (Thermo #15338030) in Opti-MEM (Thermo #31985070). For selection of stably transfected cells, cells were treated with 0.5 μg/mL puromycin (Thermo #A1113802) for 3 - 7 days. To induce expression of histone probes in transgenic cells, 0.5 µg/mL doxycycline (Cayman Chemical #14422) was supplemented in culture media. For EZH2 inhibition experiments in BT-549, 20 µM Tazemetostat (Selleck #S7128) was added to culture media for at least 5 days. All cell lines were grown at 37°C, with 5% CO2 in a humidified incubator.

### Cell Sample Preparation and Microscopy

To visualize GFP signal localization, 10×10^4^ cells were plated in a 24-well microplate 5-7 days prior to imaging. Live-cell images were acquired using a BioTek Cytation 1 (BioTek #CYT1FAV) at 20X magnification with channel settings as follows: Brightfield, CFP - ex. 445/45, em. 510/42 (Thermo #AMEP4953). To examine microenvironment-associated cues, fibronectin-coated polyacrylamide hydrogels were prepared as previously described^44^. Cells were seeded to the hydrogels at a density of ∼17,000 cells/cm^2^ and 24 hr after plating cells were treated with 0.5 µg/mL doxycycline and 10 ng/mL human TGFβ1 recombinant protein (PeproTech 100-21C-10UG) for 48 hr prior to live-cell imaging. Live-cell images were acquired using a 20X objective on a Nikon Eclipse Ti-E inverted fluorescence microscope equipped with a Photometrics CoolSNAP HQ^2^ CCD camera.

To monitor probe localization and DNA simultaneously, cells cultured on fibronectin-coated glass slides were stained with Hoechst 33342 (1:1000) for 20 minutes prior to live-cell imaging using a Leica DMI 8 confocal microscope with a 20X air objective. Z-stacks were acquired using settings as follows: GFP - 491 nm, intensity 0.2%; Hoechst 33342 - 405 nm, intensity 3.66%.

### Image Processing in FIJI

Wide field and confocal images were processed with FIJI/ ImageJ (version 2.14.0/1.54f). For machine learning model-assisted analysis, prior to image segmentation the following steps were applied: (1) Rolling ball background subtraction (radius = 5 pixels) to correct uneven illumination; (2) Contrast enhancement with a saturation fraction of 0.35%. Subnuclear dot segmentation was performed using Cellpose, a deep learning-based segmentation framework. The cyto3 pretrained model was selected, with an expected dot radius of 5 pixels. Grayscale images were used as input. Quantitative measurements were exported in CSV format, where each row corresponded to a cell and included dot count (number of segmented regions), and average dot size (total masked area divided by dot count). The python libraries used include: PyTorch, timm, tifffile, Cellpose, NumPy, matplotlib, scikit-learn, and openpyxl.

## ACKNOWLEDGEMENTS

Cell-free transcription-translation (TX-TL) mix was generously provided by M. Styczynski. Use of the Leica DMI 8 confocal microscope was generously provided by A. Sheikhi. Z. Hirani performed Gibson Assembly for the pSBtetTA-luc2GG plasmid, and J. Titus assisted with variant library plasmid preparation. This project was supported by NIH R21CA232244 (to KAH) and NSF 2414436 (to EWG).

## AUTHOR CONTRIBUTIONS

**K.A.H.** conceived the study and secured funding. **J.H.P.** optimized Golden Gate assembly, and **J.H.P.** and **K.A.F.** constructed the library clones. **P.S.** performed evolutionary analysis of Polycomb chromodomains and assisted with cloning. **K.A.F.** designed TXTL assays, optimized CHIA, screened the protein library, engineered clasp variants, constructed mammalian expression vectors, generated transgenic BT-549 cell lines, performed data analysis, and wrote the manuscript. **A.A.** and **E.W.G.** designed and performed hydrogel and confocal microscopy experiments. **I.K., J.S., and A.C.** developed and applied machine learning methods for quantitative analysis of nuclear fluorescence patterns. **K.A.H.** assisted with manuscript preparation and supervised the project. All authors reviewed and approved the final manuscript.

## DECLARATION OF INTERESTS

The authors declare no conflict of interest.

## Notes

### Competing Interest Statement

The authors have declared no competing interest.

### Summary of Updates

This version includes substantial revisions to improve the scientific rigor, organization, and biological validation of the study. The manuscript has been extensively rewritten for clarity, with revised text throughout the Introduction, Results, Discussion, and Methods sections. Figures have been redesigned and reorganized to improve data presentation and interpretation. New experiments demonstrate the utility of the engineered CBX8 variants as live cell chromatin probes in mammalian cells. These additions include expression and localization studies across multiple human cell lines, analysis of probe localization following EZH2 inhibition, and validation using breast cancer cells cultured on substrates with different mechanical properties. The revised manuscript also incorporates confocal microscopy performed on hydrogel cultured cells and a machine learning based image analysis pipeline for quantitative classification of nuclear fluorescence patterns. The engineered variant selected for downstream studies was further justified through expanded biochemical analyses, including additional comparisons of on target versus off target histone binding and ELISA titration experiments confirming measurements within the linear detection range. The supplemental information has been substantially expanded with new figures describing assay calibration, computational image analysis, additional mammalian cell imaging, and supporting biochemical data. Authorship has been updated to recognize new collaborative contributions in quantitative image analysis and confocal microscopy. The manuscript summary, figures, supplemental information, and references have also been updated to reflect these additions. Overall, this version extends the study beyond engineering and biochemical characterization to demonstrate the performance of the optimized CBX8 variant as a selective live cell probe for H3K27me3.

