## Supplemental Information for "An engineered chromatin protein with enhanced preferential binding of H3K27me3 over H3K9me3"

Figure S1. Evolutionary conservation analysis of the CBX8 PCD
Figure S2. CBX8 PCD variant library and Golden Gate assembly validation
Figure S3. CHIA ELISA data by CBX8 variant sub-group
Figure S4. Underlying data for results shown in Figure 4C
Figure S5. Titration experiments to determine the linear range of CHIA signals
Figure S6. Expression of the CBX8.IEI-GFP fusion in cultured human cell lines
Figure S7. Computational classification of nuclei in HBD-GFP-expressing breast cancer cells with and without EZH2 inhibition
Figure S8. Microenvironment-associated cues modulate CBX8.IEI-GFP nuclear patterning

Table S1. Primers used to generate donors for Golden Gate assembly

Appendix A1. DNA sequences encoding protein domains used to assemble the fusion proteins
Appendix A2. List of codon substitutions for select CBX8 variants
Appendix A3. Computational pipeline for the HBD-GFP-expressing nucleus classifier

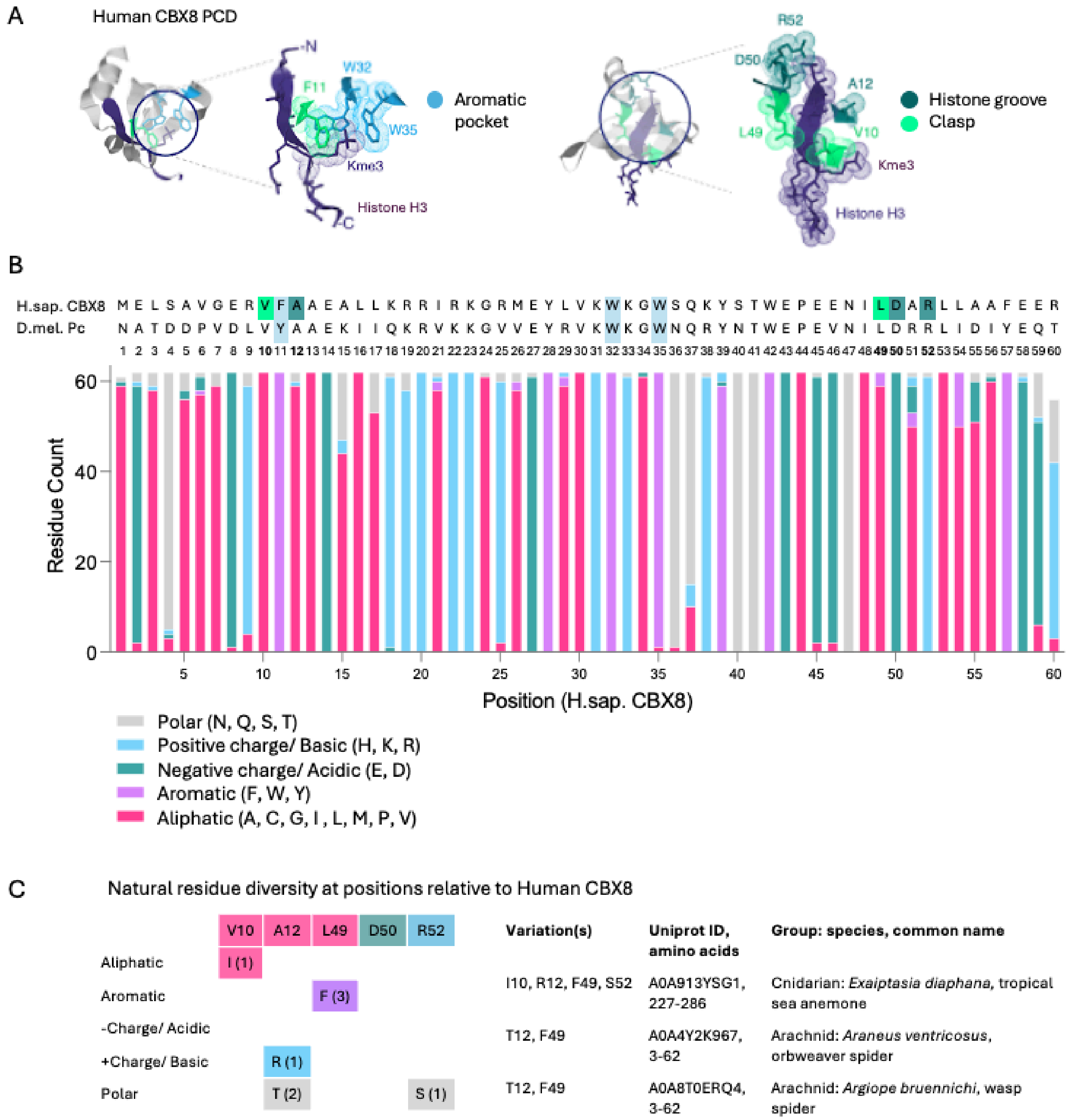


**Figure S1. Evolutionary conservation analysis of the CBX8 PCD.** A. Structure of the PCD domain from human CBX8 (RCSB PDB 3I91). Target sites for substitutions in the CBX8 variant library are highlighted green and teal. B. Sequences for human CBX8 (M1-R60) and Drosophila Pc (N16-T75) are shown at the top. Conserved aromatic cage residues F/Y, W, and W are highlighted in pale blue. Orthologous sequences from other species were retrieved by querying the 60 amino-acid human CBX8 PCD (M1 - R60) in NCBI-BLAST (blastp). We confirmed the identities of these proteins as CBX8 rather than closely related orthologs (i.e. CBX2, 4, 6, 7) by manual cross-check with entries in Uniprot. Additional CBX8 sequences in related species were searched manually in Uniprot. Ungapped sequence comparisons were performed manually by using the first two aromatic cage residues as an anchor point. After alignment, occurrences of alternative residues were tallied (stacked bar graph). C. Natural variations at the five target sites that were artificially mutated in the synthetic CBX8 variant library.

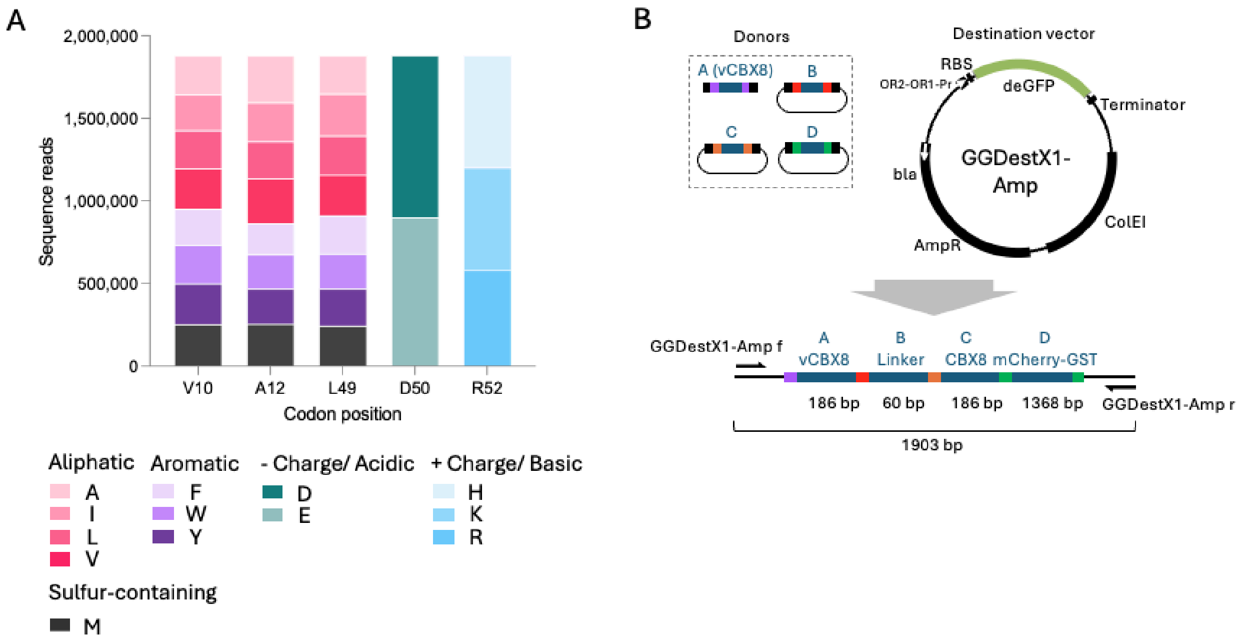


**Figure S2. CBX8 PCD variant (vCBX8) library and Golden gate assembly validation.**

A. PCD variants flanked by Golden Gate adaptors for position A were generated by Genscript as a combinatorial library with the following codon substitutions:

V10 (GTG) to A (GCC), I (ATC), L (CTG), M (ATG), F (TTC), W (TGG), or Y(TAC).

A12 (GCG) to V (GTG), I (ATC), L (CTG), M (ATG), F (TTC), W (TGG), or Y (TAC).

L49 (CTG) to V (GTG), A (GCC), I (ATC), M (ATG), F (TTC), W (TGG), or Y (TAC).

D50 (GAT) to E (GAG).

R52 (CGC) to H (CAC), or K (AAG).

Deep sequencing was used to verify that codon representation was evenly distributed. B. In the fusion constructs, vCBX8 fragments were placed at the first Golden Gate assembly position “A.” The tandem PCD domains connected by a flexible linker, along with a fluorescent mCherry reporter and GST tag, were assembled into expression vector GGDestX1-Amp. Individual variants were isolated by transforming *E. coli* DH5α-Turbo with the Golden Gate reaction product and picking single colonies from LB ampicillin agar plates.

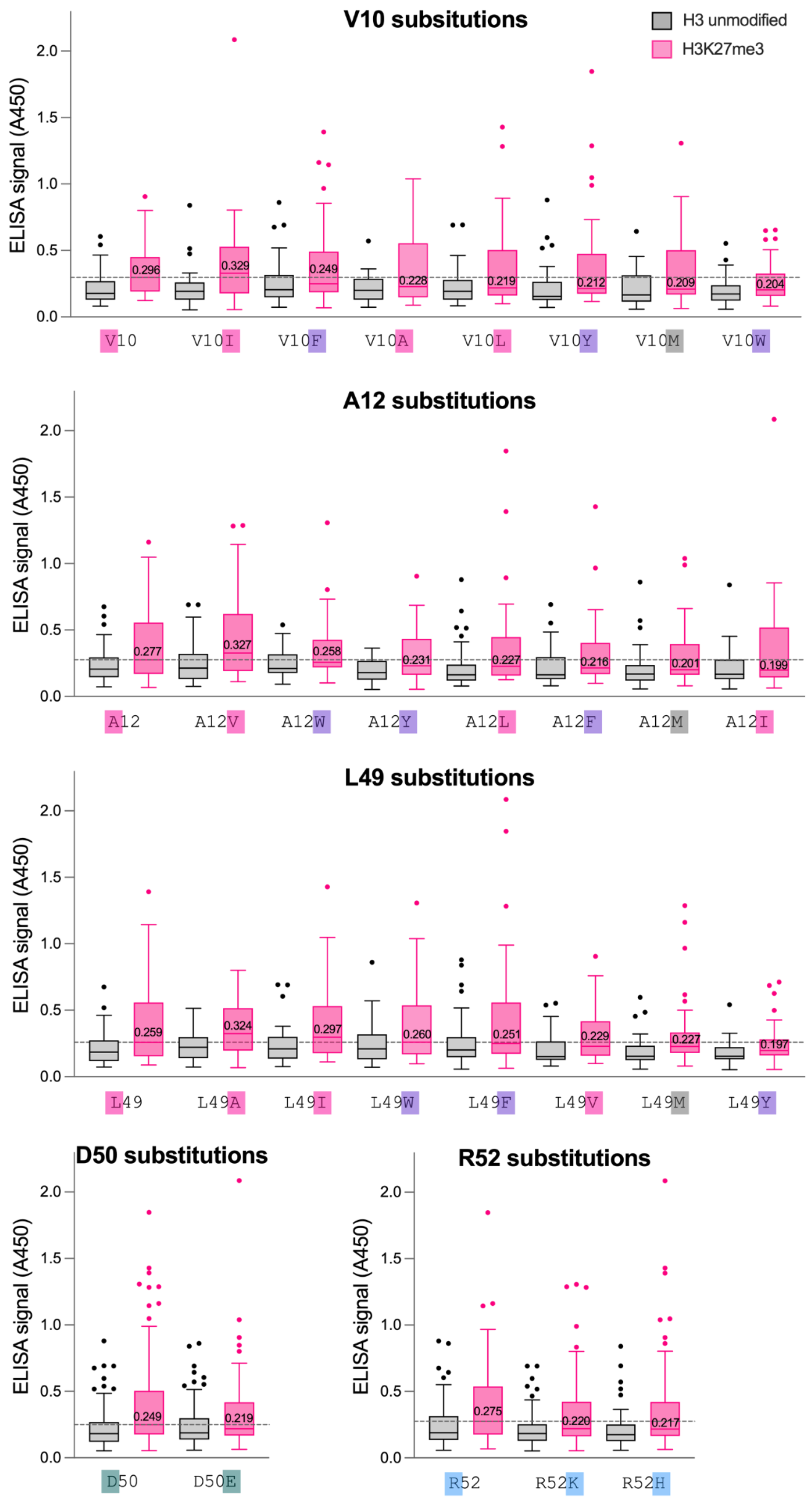


**Figure S3. CHIA ELISA data by CBX8 variant sub-group.**

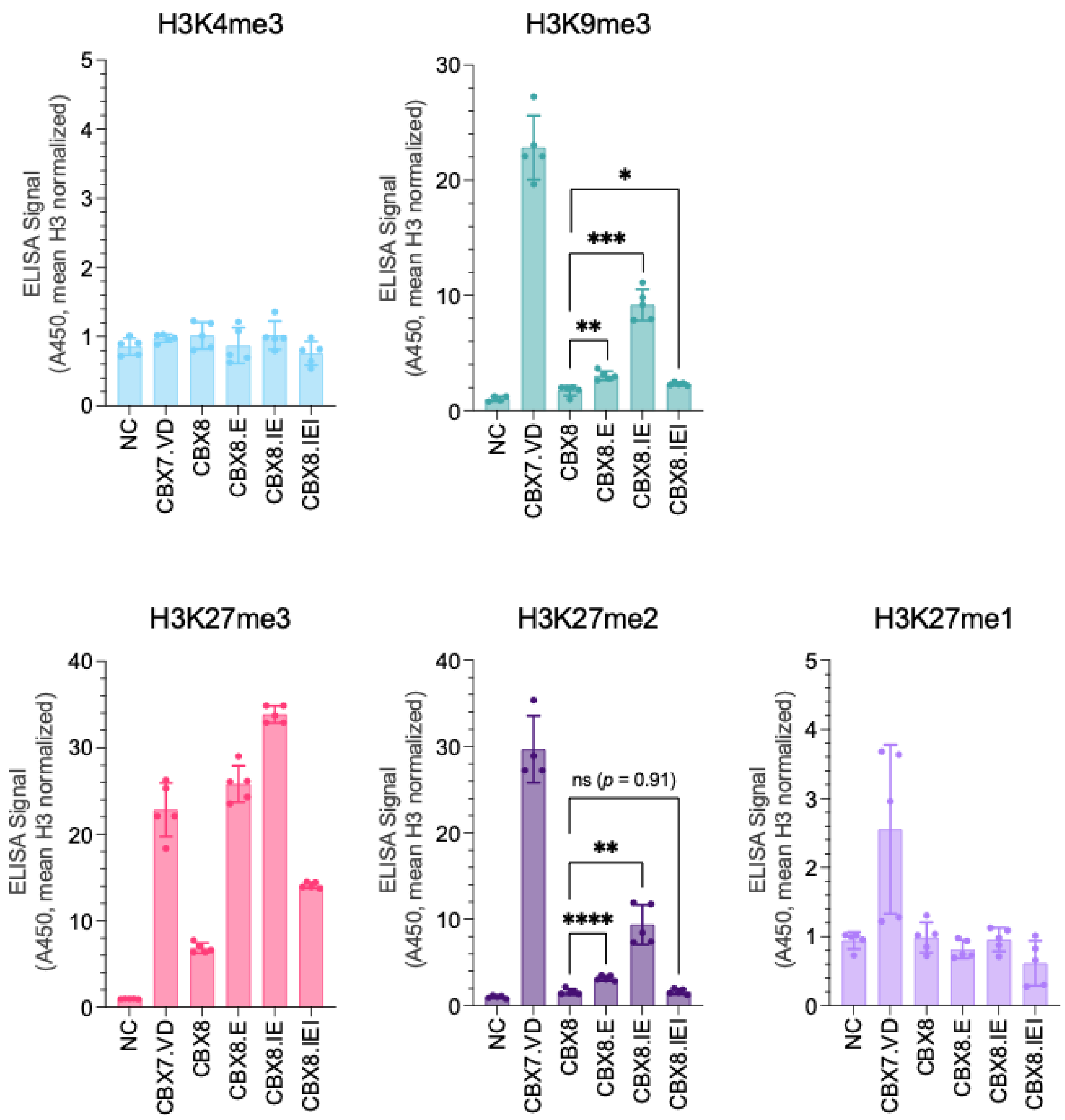


**Figure S4. Underlying data for results shown in Figure 4C.** Data are plotted from the measurements underlying the data shown in the heatmap in Figure 4C to facilitate direct comparison of off-target binding between H3K27me3 reader variants. Dots represent individual technical replicates, bars indicate the mean, and error bars denote standard deviation. These results motivated the selection of CBX8.IEI for subsequent mammalian cell expression studies. NC = negative control Welch's t-test, two-tailed, unpaired: ns; * $p < 0.05$; ** $p < 0.01$; *** $p < 0.001$.

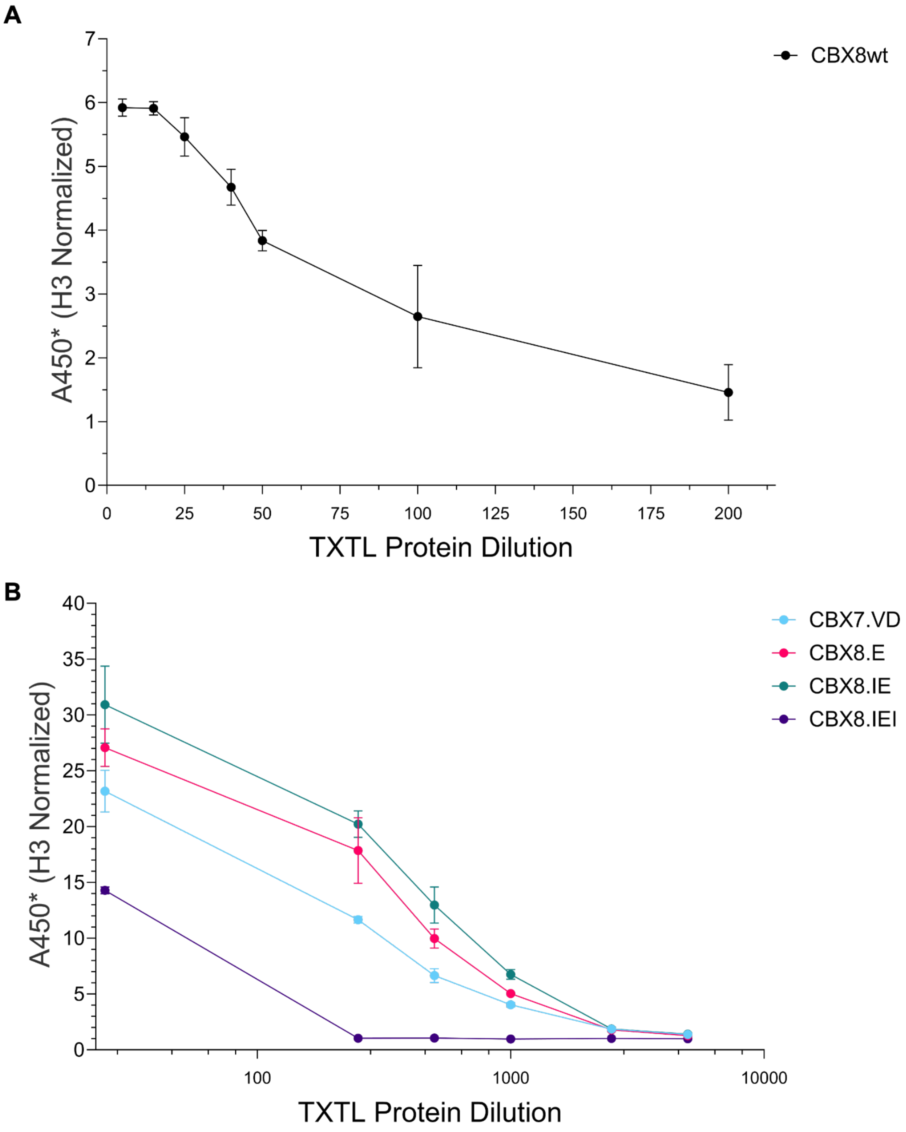


**Figure S5. Titration experiments to determine the linear range of CHIA signals.** A. To determine the linear range of the ELISA signal for the CHIA screen, the TXTL product for homotypic CBX8 (w.t.) was tested at dilutions from 1:5 to 1:200 B. TXTL products for homotypic CBX7.VD, CBX8.E, CBX8.IE, and CBX8.IEI showed saturating ELISA signal (A450 > 4.0) when applied at the standard dilution (1:25) in some of the replicates. To verify the relative signals (CBX8.IEI > CBX8.IE > CBX7.VD > CBXB.IEI) reported in Fig. 4, TXTL products were serially diluted (1:25 - 1:5000) and used for ELISA. Dots = A450 from 3 technical replicate wells, error bars = standard deviation.

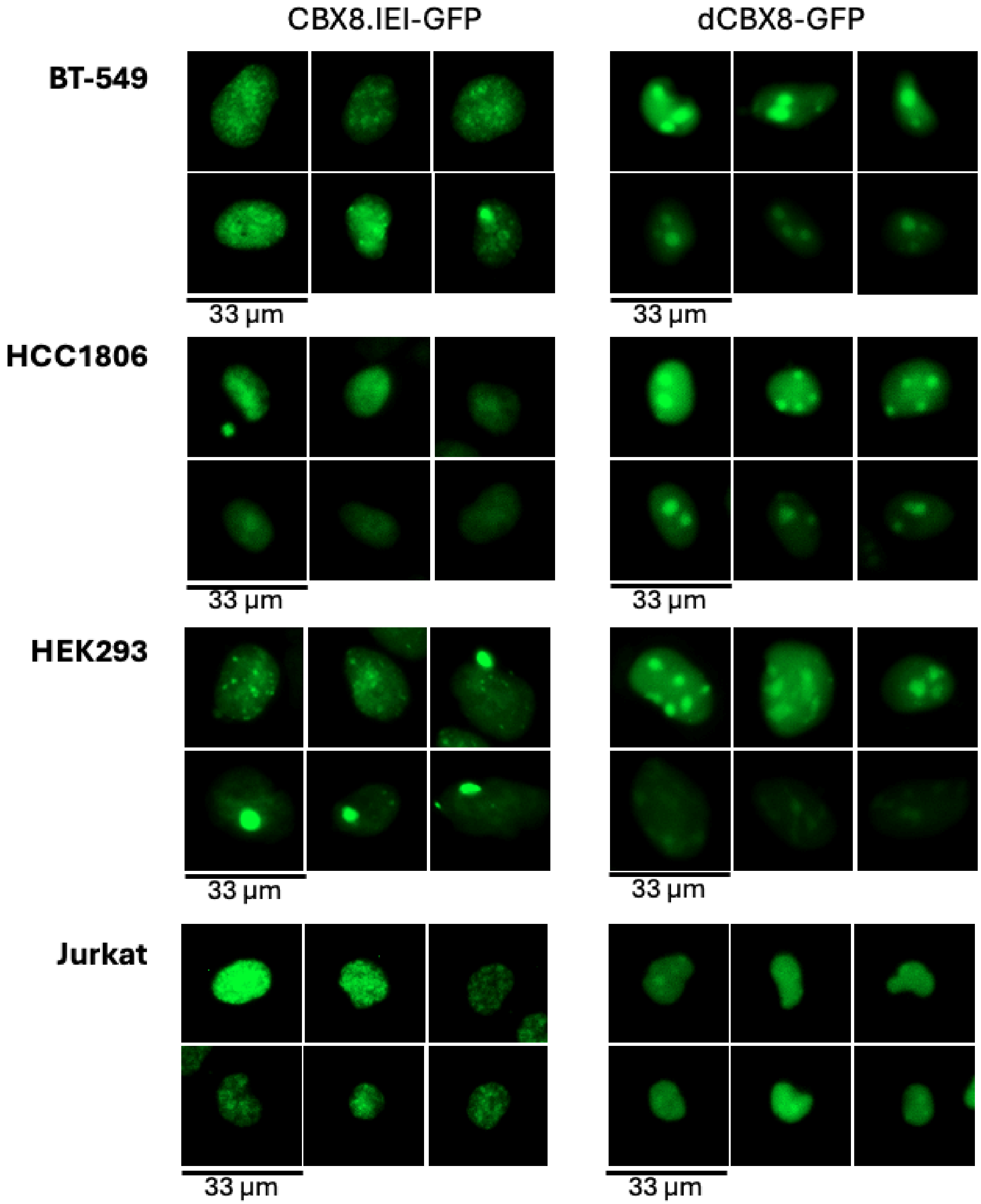


**Figure S6. Expression of the CBX8.IEI-GFP and dCBX8-GFP fusions in cultured human cell lines.** Representative fluorescence microscopy images of doxycycline-induced CBX8.IEI-GFP and dCBX8-GFP in BT-549, HCC1806, HEK293, and Jurkat cells. Images were selected to illustrate the range of nuclear localization patterns observed across cell types, including diffuse, compartmentalized, and punctate distributions. These examples complement the representative images shown in Figure 5B.

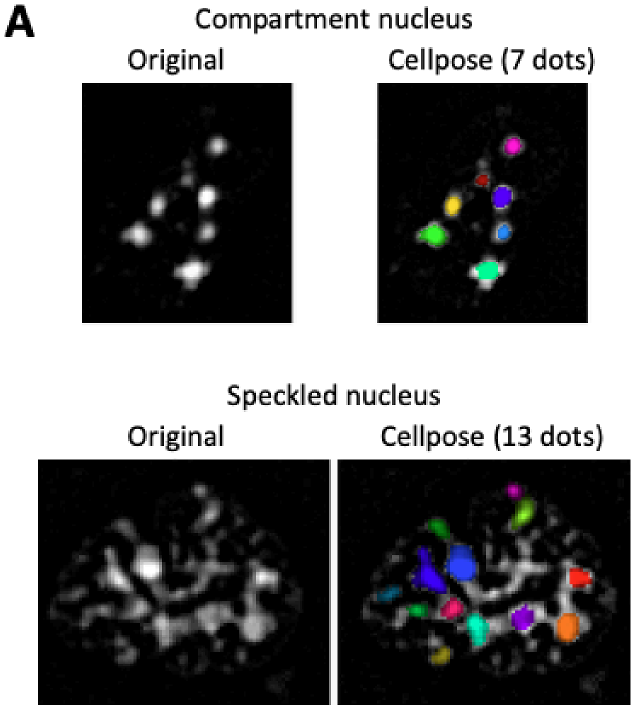


**B**

| | Precision | Recall |
|---|---|---|
| Compartment | 0.857 | 0.750 |
| Speckled | 0.962 | 0.758 |
| Unclassified | 0.000 | 0.000 |

**Figure S7. Computational classification of nuclei in HBD-GFP-expressing breast cancer cells with and without EZH2 inhibition.** A. Representative examples of nuclei classified as “compartment” or “speckled.” Left, original fluorescence images. Right, output from Cellpose-based nuclear segmentation and punctae detection, with individual dots pseudo-colored for visualization and counted per nucleus. B. Performance metrics of the machine learning classifier trained to distinguish compartment and speckled phenotypes. Precision and recall values are shown for each class. “Unclassified” nuclei reflect cases not confidently assigned to either phenotype.

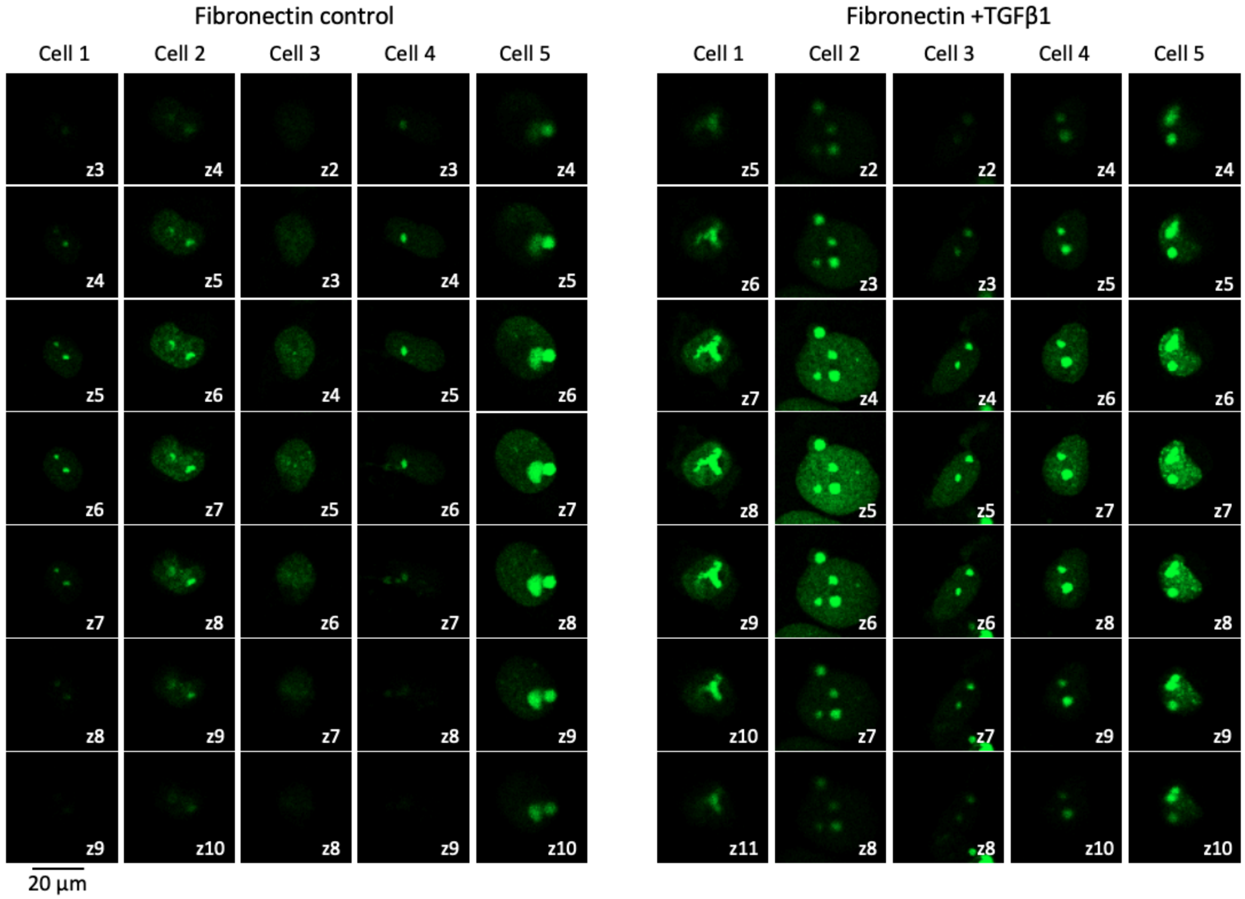


**Figure S8. Microenvironment-associated cues modulate CBX8.IEI-GFP nuclear patterning.** Representative confocal z-plane series of BT-549 cells expressing CBX8.IEI-GFP cultured on fibronectin-coated glass under control conditions (left) or following TGFβ1 stimulation (right). Individual nuclei (Cells 1 - 5) are shown across sequential optical sections, with z-plane indices indicated. Images are representative of multiple fields acquired under identical imaging settings for each construct (Nikon Eclipse Ti-E microscope; 20x objective; FITC filter; 20 ms exposure).

**Table S1.** Primers used to generate donors for Golden Gate assembly. Sequences of only the template-binding regions are shown. GG Module A, B, C, or D-specific 5’ extensions (lowercase) were added to the sequences shown below to generate full-length primers, as described in Haynes & Priode 2023[1]. In some cases a start codon (atg), or stop codon (tag, r.c. cta) was included in the primer extension (underlined text).

| Protein domain | GG Module(s) | Forward (fwd) and reverse (rev) primer sequences (5’...) |
|---|---|---|
| CBX8 and variants | A | Fwd ttgaagacctggagATGGAGCTTTCAGCGGTGG<br>Rev ttgaagaccccagaTCTTTCCCTTTCCTCAAAGGC |
| | C | Fwd ttgaagacctccatccATGGAGCTTTCAGCGGTGG<br>Rev ttgaagacccgggcTCTTTCCCTTTCCTCAAAGGC |
| BAH | A | Fwd ttgaagacctggagGCCAAAGAGCTCTCCCGGAGGCA<br>Rev ttgaagaccccagaGATGGGCACGCCATCAGCCGTC |
| CBX7.VD | A | Fwd ttgaagacctggagATGGAACTGAGCGCG<br>Rev ttgaagaccccagaTTTCGGGCCGCGTTT |
| | C | Fwd ttgaagacctccatccATGGAACTGAGCGCG<br>Rev ttgaagacccgggcTTTCGGGCCGCGTTT |
| CBX1 | A | Fwd ttgaagacctggagatgGAAGAAGAAGAAGAAGAATATGTGG<br>Rev ttgaagaccccagaGGTTTTCTGAGATTGTAGAAATTCG |
| | C | Fwd ttgaagacctccatccatgGAAGAAGAAGAAGAAGAATATGTGG<br>Rev ttgaagacccgggcGGTTTTCTGAGATTGTAGAAATTCG |
| CBX1eR | A | Fwd ttgaagacctggagatgGAAGAAGAGGAAGAGGAATATGTG<br>Rev ttgaagaccccagaTGTTTTCTGTGACTGTAGAAACTC |
| | C | Fwd ttgaagacctccatccatgGAAGAAGAGGAAGAGGAATATGTG<br>Rev ttgaagacccgggcTGTTTTCTGTGACTGTAGAAACTC |
| Linker 4X[GGGGS] | B | Fwd ttgaagaccttctggaGGCGGTGGCGGATCT<br>Rev ttgaagacccatggTGAACCACCTCCTCCAGATC |
| mCherry-GST | D | Fwd ttgaagacctgcccctGTGAGCAAGGGCGAGGAGGATAACATGG<br>Rev ttgaagaccctagtATCCGATTTTGGAGGATGGTCGCCACC |
| eGFP-NLS-stop | D | Fwd ttgaagacctgcccctATGGTGAGCAAGGGCG<br>Rev ttgaagaccctagtctaTACCTTGCGCTTTTTCTTGGG |

**Appendix A1. DNA sequences encoding protein domains used to assemble the fusion proteins.** Sequences are shown in FASTA format.

```
>CBX8
atggagctttcagcggtgggggagcgggtgttcgcggccgaagccctcctgaagcggcgcatacggaaaggacgcatgg
aatacctcgtgaaatggaagggatggtcgcagaagtacagcacatgggaaccggaggaaaacatcctggatgctcgctt
gctcgcagcctttgaggaaagggaaaga

>BAH
gccaaagagctctcccggaggcagcggccgccctccgtggaaaaccggccaaagatctcagccttcctgcccgcccggc
agctctggaagtggtcggggaatcccacacagcggcgtggcatgaaggggaaggcccggaagctgttctacaaggccat
cgtgcggggcgaggagaccctgcgtgtcggggactgtgccgtAttcctgtcagctgggcggcccaacctcccctacatc
ggccgcatcgagagcatgtgggagtcgtggggcagcaacatggtggtcaaggtcaagtggttctaccaccctgaggaga
ccaagctgggcaagaggcagtgcgacggcaagaatgcgctgtaccagtcctgccacgaggatgagaacgacgtgcagac
```

```
catctcccacaagtgccaggtcgtggcgcgcgagcagtatgagcagatggcccggagccgcaagtgccaggaccggcag
gacctctactacctggcgggcacctacgaccccaccaccgggcgcctggtgacggctgatggcgtgcccatc

>CBX7.VD
atggaactgagcgcgattggcgaagatgtgtttgcggtggaaagcattcgcaaaaaacgcgtgcgcaaaggcaaagtgg
aatatctggtgaaatgggaaggctggccgccgaaatatagcacctgggaaccggaagaacatattctggatccgcgcct
ggtgatggcgtatgaagaaaaagaagaacgcgatcgcgcgagcggctatcgcaaacgcggcccgaaa

>CBX1 wildtype
gaagaaGAGgaaGAGgaatatgtggttgaaaaagtgttagatcgccgtgtggttaaaggtaaagttgaatatctgctga
aatggaaaggctttagtgatgaagataatacctgggaaccggaagaaaatctggattgtccggatctgattgccgaatt
tctacaatctcagaaaacc

>CBX1eR
Gaagaagaggaagaggaatatgtggtggaaaaagttcttgatcggcgagttgtcaagggcaaggtggaatatcttctaa
agtgggccggtttctcagatgaggacaacacttgggagccagaagagaatctgttttgccctgaccttattgctgagtt
tctacagtcacagaaaaca

>Linker 4X[GGGGS]
ggcggtggcggatctggaggtggtggttcaggcggaggcggatctggaggaggtggttca

>mCherry-GST mCherry_1-705 ACTAGAscar_706-711 GST_712-1368
gtgagcaagggcgaggaggataacatggccatcatcaaggagttcatgcgcttcaaggtgcacatggagggctccgtga
acggccacgagttcgagatcgagggcgagggcgagggccgcccctacgagggcacccagaccgccaagctgaaggtgac
caagggtggcccctgcccttcgcctgggacatcctgtcccctcagttcatgtacggctccaaggcctacgtgaagcac
cccgccgacatccccgactacttgaagctgtccttccccgagggcttcaagtgggagcgcgtgatgaacttcgaggacg
gcggcgtggtgaccgtgacccaggactcctccttgcaggacggcgagttcatctacaaggtgaagctgcgcggcaccaa
cttcccctccgacggccccgtaatgcagaagaaaaccatgggctgggaggcctcctccgagcggatgtaccccgaggac
ggcgccctgaagggcgagatcaagcagaggctgaagctgaaggacggcggccactacgacgctgaggtcaagaccacct
acaaggccaagaagcccgtgcagctacccggcgcctacaacgtcaacatcaagttggacatcacctcccacaacgagga
ctacaccatcgtggaacagtacgaacgcgccgagggccgccactccaccggcggcatggacgagctgtacaagACTAGA
TCCCCTATACTAGGTTATTGGAAAATTAAGGGCCTTGTGCAACCCACTCGACTTCTTTTGGAATATCTTGAAGAAAAAT
ATGAAGAGCATTTGTATGAGCGCGATGAAGGTGATAAATGGCGAAACAAAAAGTTTGAATTGGGTTTGGAGTTTCCCAA
TCTTCCTTATTATATTGATGGTGATGTTAAATTAACACAGTCTATGGCCATCATACGTTATATAGCTGACAAGCACAAC
ATGTTGGGTGGTTGTCCAAAAGAGCGTGCAGAGATTTCAATGCTTGAAGGAGCGGTTTTGGATATTAGATACGGTGTTT
CGAGAATTGCATATAGTAAAGACTTTGAAACTCTCAAAGTTGATTTTCTTAGCAAGCTACCTGAAATGCTGAAAATGTT
CGAAGATCGTTTATGTCATAAAACATATTTAAATGGTGATCATGTAACCCATCCTGACTTCATGTTGTATGACGCTCTT
GATGTTGTTTTATACATGGACCCAATGTGCCTGGATGCGTTCCCAAAATTAGTTTGTTTTAAAAAACGTATTGAAGCTA
TCCCACAAATTGATAAGTACTTGAAATCCAGCAAGTATATAGCATGGCCTTTGCAGGGCTGGCAAGCCACGTTTGGTGG
TGGCGACCATCCTCCAAAATCGGAT

>eGFP-NLS eGFP_1-717 SV40NLS_718-739
atggtgagcaagggcgaggagctgttcaccggggtggtgcccatcctggtcgagctggacggcgacgtaaacggccaca
agttcagcgtgtccggcgagggcgagggcgatgccacctacggcaagctgaccctgaagttcatctgcaccaccggcaa
gctgcccgtgccctggcccaccctcgtgaccaccctgacctacggcgtgcagtgcttcagccgctaccccgaccacatg
aagcagcacgacttcttcaagtccgccatgcccgaaggctacgtccaggagcgcaccatcttcttcaaggacgacggca
actacaagacccgcgccgaggtgaagttcgagggcgacaccctggtgaaccgcatcgagctgaagggcatcgacttcaa
ggaggacggcaacatcctggggcacaagctggagtacaactacaacagccacaacgtctatatcatggccgacaagcag
aagaacggcatcaaggtgaacttcaagatccgccacaacatcgaggacggcagcgtgcagctcgccgaccactaccagc
agaacacccccatcggcgacggccccgtgctgctgcccgacaaccactacctgagcacccagtccgccctgagcaaaga
ccccaacgagaagcgcgatcacatggtcctgctggagttcgtgaccgccgccgggatcactctcggcatggacgagctg
tacaagcccaagaaaaagcgcaaggta
```

**Appendix A2. List of codon substitutions for select CBX8 variants.** Substitutions were generated in the CBX8 sequence from Appendix A1.

CBX8 K33: K33 (AAG) to E (CGC)

CBX8 V10I K33: V10 (GTG) to I (ATC), (AAG) to E (CGC)

CBX8 V10I K33 L49I: V10 (GTG) to I (ATC), (AAG) to E (CGC), L49 (CTG) to I (ATC)

**Appendix A3. Computational pipeline for the HBD-GFP-expressing nucleus classifier.**

*Overview*

The computational pipeline consisted of four stages:

1. Few-shot image classification using a self-supervised Vision Transformer,
2. Image processing and normalization (see main Methods: Image Processing in FIJI),
3. Automated cell segmentation and dot detection using Cellpose, and
4. Quantitative feature extraction and analysis.

Except for FIJI preprocessing, which was automated using FIJI macros in its native environment, the entire workflow was implemented in Jupyter Notebook to ensure reproducibility. All analyses were performed in Python (version 3.10).

*Rationale for Model Selection*

The DINOv2 (Distillation with No Labels v2) model developed by Meta AI was selected for the classification of fluorescence patterns in the nucleus images. DINOv2 is a self-supervised Vision Transformer that produces high-quality image embeddings without task-specific fine-tuning. Conventional convolutional neural networks typically require large labeled datasets for effective training. Because our dataset contained a relatively small number of labeled fluorescence images, a model capable of generating robust feature representations under limited supervision was preferred. DINOv2 satisfied this requirement.

*Training Data and Classification Strategy*

Cell nuclei were randomly cropped and manually classified into two morphological categories: compartment or speckle. Labeled images were organized into class-specific directories. All labeled images were used as support examples from which DINOv2 embeddings were extracted. A total of 152 images were labeled, comprising 76 compartment and 76 speckle samples.

To reduce inter-image intensity variability, per-image contrast normalization was performed using an interquartile range (IQR)-based approach:

$$Normalized\ intensity = \frac{(I - median)}{3 * IQR} + 0.5$$

where IQR = Q3 − Q1 of the image intensity distribution.

This normalization emphasizes relative intensity structure rather than absolute brightness. Because DINOv2 requires three-channel input, the grayscale image was replicated across three channels. Channel-wise normalization was then applied using a mean of 0.5 and a standard deviation of 0.25 to match the distribution expected by the pretrained model. The resulting 3-channel tensor was then submitted to the DINOv2 backbone for embedding extraction.

*Embedding Representation and Classification*

For each image, DINOv2 produced a 384-dimensional embedding vector derived from the CLS token representation. The model partitions the 518 x 518 image into 14 x 14 pixel patches, encodes each patch, and aggregates global information via the CLS token, which summarizes the entire image. Each embedding vector was L2-normalized to unit length to enable cosine similarity comparisons. Embeddings were extracted for all support (labeled) and query (unlabeled) images. Classification of query images was performed using k-nearest neighbors (k-NN) with:

- k = 5
- Cosine similarity as the distance metric

For each query image, the five closest support embeddings were identified. Weighted voting was performed using cosine similarity scores to produce the final class prediction.

The model was initially tested with 152 labeled images with 76 compartments and 76 speckle to assess accuracy of the model before applying to the larger set of images that are unlabeled. 80% (n=122) of the images are used as a query to train the DINOv2 model while 20% (n=30) were used as a query to test the accuracy and precision of the classification model. The model achieved a weighted f1-score of 0.866 and an overall accuracy of 86.7%. Per-class F1 scores were 0.857 for compartment images and 0.875 for speckle images. The confusion matrix indicated that there were 12/15 compartments, and 14/15 speckle images were correctly classified.

To assess the generalizability of the DINOv2-based classifier, 6-fold stratified cross-validation was performed on the 152 labeled images (76 compartment and 76 speckle). Stratification ensured balanced class representation within each fold, resulting in approximately 126 support images and 12–13 test images per class in each validation split.

Because the DINOv2 backbone was frozen and used solely as a feature extractor, embeddings for all 152 images were computed once and reused across folds to avoid redundant computation. For each fold, classification was performed using a k-nearest neighbors (k-NN) classifier with k = 5 and cosine similarity as the distance metric.

Model performance was evaluated using overall accuracy, per-class precision, recall, and F1-score, as well as weighted-average precision, recall, and F1-score:
The model achieved a mean accuracy of 0.846 ± 0.066 across folds, with weighted precision, recall, and F1-scores of 0.855 ± 0.069, 0.843 ± 0.066, and 0.842 ± 0.066, respectively.
Per-class F1-scores were 0.830 ± 0.070 (compartment) and 0.853 ± 0.064 (speckle).

*Confidence Thresholding*

To reduce ambiguous predictions, a confidence-based filtering strategy was applied. For each query image, weighted votes from the five nearest neighbors were aggregated by class.
An image was classified as unsure if either of the following conditions was met:

1. The confidence score (defined as the weighted vote of the top class divided by total votes) was < 0.60.
2. The margin between the top two class scores was < 0.15.

Images classified as unsure were excluded from downstream analysis to improve reliability.
Confident predictions were automatically saved in class-specific directories.
